# Developmental stability and segregation of Theory of Mind and Pain networks carry distinct temporal signatures during naturalistic viewing

**DOI:** 10.1101/2023.08.09.552564

**Authors:** Km Bhavna, Niniva Ghosh, Romi Banerjee, Dipanjan Roy

**Affiliations:** Department of Computer Science and Engineering, Indian Institute of Technology, Jodhpur, Rajasthan, India; School of Artificial Intelligence Data Science, Centre for Brain Science Application, Indian Institute of Technology, Jodhpur, Rajasthan, India

**Keywords:** Theory of Mind, Pain Networks, Angular distance, Dynamic functional connectivity, Inter Subject Correlations

## Abstract

Temporally stable large-scale functional brain connectivity among distributed brain regions is crucial during brain development. Recently, many studies highlighted an association between temporal dynamics during development and their alterations across various time scales. However, systematic characterization of temporal stability patterns of brain networks that represent the bodies and minds of others in children remains unexplored. To address this, we apply an unsupervised approach to reduce high-dimensional dynamic functional connectivity (dFC) features via low-dimensional patterns and characterize temporal stability using quantitative metrics across neurodevelopment. This study characterizes the development of temporal stability of the Theory of Mind (ToM) and Pain networks to address the functional maturation of these networks. The dataset used for this investigation comprised 155 subjects (children (n=122, 3–12 years) and adults (n=33)) watching engaging movie clips while undergoing fMRI data acquisition. The movie clips highlighted cartoon characters and their bodily sensations (often pain) and mental states (beliefs, desires, emotions) of others, activating ToM and Pain network regions of young children. Our findings demonstrate that ToM and pain networks display distinct temporal stability patterns by age 3 years. Finally, the temporal stability and specialization of the two functional networks increase with age and predict ToM behavior.

## 1. Introduction

THEORY-OF-MIND (TOM) is an ability to understand other’s mental states and was highlighted in 1970 by Premack and Woodruff [1]. This ability allow one to comprehend other people’s aims, belief, ambitions, emotions, and mentalization of concepts that differs from one’s own [2]. Recent studies focus on the early development of Theory-of-mind in children trying to understand the development of ToM ability beyond the preschool age group, and its association with middle childhood and early adolescence and individual differences [3]. According to the extant literature, children develop the ability to understand faith and desire and predict another person’s actions as seen in false belief task paradigms by age 5 [3], [4], [5], [6]. Previous studies have suggested that children’s ability to anticipate or justify another person’s actions based on false beliefs depends on the development of understanding concepts during ToM development that happens around the age of four years [4]. Hence, children’s performance in explicit false-belief tasks could index an important milestone in understanding concepts during ToM development. During maturation, there is a dramatic alteration in the representation of others’ internal states, providing critical insight into children’s social cognition ability development. Therefore, investigating the early age group allows us to examine mentalization and re-examine the literature about the child’s ToM based on predictions based on social brain regions of children 3-12 years during naturalistic movie-watching tasks. The human brain is a collection of massive functional modules that become more distinct during development from childhood to adolescence, i.e., connectivity within modules increases as we grow from childhood to adolescence, and connectivity between modules decreases [7], [8], [9], [10], [11]. According to this view, this could also reflect concurrent development in other domain-general brain regions such as the Default mode Network a cluster of brain regions implicated in self-related processing [12]. Therefore, previous studies have focused on functional connectivity measures within and between ToM and DMN to address children’s early developmental differences and functional specialization in 3-12 years and relate that to performance in explicit false-belief reasoning tasks. Moreover, previous studies have reported a clear difference between brain areas responding preferentially to internal states of others’ bodies (like hunger, pain) versus internal states (like beliefs, emotions, and desires) of others’ minds suggesting a division of labor and early segregation into functionally specialized brain regions [12], [4]. Previous studies investigated the differences in temporal stability of functional architecture in the resting states of patients with neurological disorders and healthy controls and examined the effects of various activities [11], [13], [14], [15], [16]. The studies also demonstrated neurological disorders (e.g., ADHD, schizophrenia, and autism spectrum disorder) specific variable alterations in the functional architecture of the default mode network (DMN), visual areas of the brain, and subcortical regions of the brain [17], [11], [13], [14], [15]. Nonetheless, quantitative characterization of the temporal stability of these functionally specialized brain regions across neurodevelopment remains completely unexplored. Therefore, there is a genuine knowledge gap in neuroimaging studies concerning the early stages of development in ToM ability.

Existing neuroimaging studies further suggest during ToM-related tasks, neural activation has been found predominantly in the Bilateral Temporoparietal Junction (Left TPJ and Right TPJ), Medial Prefrontal Cortex, and Posterior Cingulate Cortex using fMRI data; however, this leaves a genuine gap in understanding what precisely the age range in neurodevelopment when temporal stability of these regions is being attended [12]. Moreover, there are no existing studies that quantify whether the temporal stability of social brain regions of ToM and Pain (sensory) networks carry distinct or overlapping signatures during naturalistic stimulus processing in children during development and whether the pattern of temporal stability indicates the successful development of ToM reasoning ability thus far. Secondly, whether the temporal stability patterns of social brain regions of ToM in 3-12 years could predict the successful performers in False Belief reasoning tasks, Finally, whether the behavioral performances can also be predicted based on inter-subject correlations (ISCs) and their association with temporal stability of ToM and Pain networks [18], [4] remains least understood. To address the above questions, first, we analyze high-dimensional dynamic functional connectivity (dFC) via low-dimensional representations. Thereafter, estimating temporal stability using quantitative metrics in 122 children and a reference group of 33 adults. Temporal stability was estimated from fMRI data while participants were viewing a short, animated movie that included events evoking the mental states and physical bodily sensations of the clip characters. A recent study has validated this movie which activates ToM and the pain brain regions in adults [4].

Subsequently, we use Angular and Mahalanobis distance to identify the ToM and Pain network temporal stability when movie stimuli/clips contain internal activation of mental and physical states activating brain networks at specific epochs of time points. We tested our hypothesis that spontaneous processing of others’ mental states within ToM brain networks might exhibit similar hyper-connectivity patterns (high inter-subject correlations) in children who pass and a and hypo connectivity (lower inter-subject correlations) patterns in domain-specific regions for participants who fail explicit false-belief tasks. Third, successful performers and their temporal stability associated with development could shed new insight into the ongoing conceptual development of ToM, which begins early in development—and continues till early adolescence displaying high temporal instability at an early age and reaching stabilization as ToM conceptual network continues to develop further. We have conducted the following analysis to address core questions regarding the temporal stability of functional brain networks (ToM and Pain) involved in representing internal mental and physical states.

The brain regions associated with ToM and pain networks have been selected based on a previous study that reported 12 brain regions (Refer to Table 1) [19], [20]. ToM brain regions include bilateral temporoparietal junction, precuneus, and dorso-, middle-, and ventromedial prefrontal cortex. The pain network comprises brain regions recruited when perceiving the physical pain and bodily sensations of others: bilateral medial frontal gyrus, insular cortex, secondary sensory cortex, and dorsal anterior middle cingulate cortex. As in previous studies, we collapse these brain regions across specific functions and use ToM and pain networks as regions generally recruited for reasoning about others’ internal mental and physical states. The contribution of the current study is as follows: **A)** We used the dynamic functional connectivity (dFC) to quantify temporal stability using our previous approach [15] of functional brain networks in 3-12 years children. Subsequently, we estimated dominant dFC subspaces of ToM and Pain networks to quantify their segregation and differences at early age. **B)** To further capture ToM network temporal stability, we quantified differences between the dominant dFC subspaces using (i) Angular Distance and (ii) Mahalanobis Distance. Our results indicate that ToM and Pain networks are achieving considerable temporal stability by the age of 5 yrs. **C)** Finally, we have tested whether the temporal stability of functional networks could predict whether a participant could pass, fail or give an inconsistent response in a false-belief reasoning task carried out by the 3-12 years old participants outside the scanner. Finally, we have empirically measured Inter-Subject Connectivity (ISCs) within ToM and Pain networks of participants ([21], [22], [23]) to test our hypothesis that both temporal stability and ISC of functional brain networks were related and could track reliably developmental differences in ToM and Pain networks of participants. Our finding grounded on derived temporal stability differences in functional brain networks during developmental age may aid in developing avenues to address questions related to distinct neural response to others’ minds and bodies that is present at a very early age and throws fundamental insights into the functional maturation of ToM brain networks.

**TABLE I:** Table shows selected ToM and pain brain regions and corresponding MNI-coordinated for extracting time-series signal.

| ToM Regions |  |  | Pain Regions |  |  |
| --- | --- | --- | --- | --- | --- |
| Sr. No. | ROIs | MNI-Coordinates (X,Y,Z) | Sr. No. | ROIs | MNI-Coordinates (X,Y,Z) |
| 1 | Posterior Cingulate Cortex (PCC) | 0, -52, 18 | 1 | Right Middle Frontal Gyrus (RMFC) | 36, 38, 40 |
| 2 | Left Temporoparietal Junction (LTPJ) | -46, -68, 32 | 2 | Left Middle Frontal Gyrus (LMFC) | -36, 38, 40 |
| 3 | Right Temporoparietal Junction (RTPJ) | 46, -68, 32 | 3 | Left Interior Insula (LII) | -40, 22, 0 |
| 4 | Ventromedial Prefrontal Cortex (vmPFC) | 4, 48, -4 | 4 | Right Interior Insula (RII) | 39, 23, -4 |
| 5 | Precuneus | 0, -49, 40 | 5 | Left Secondary Sensory Cortex (LSSC) | -39, -15, 18 |
| 6 | Dorsomedial Prefrontal Cortex (dmPFC) | -10, 58, 24 | 6 | Right Secondary Sensory Cortex (RSSC) | 39, -15, 18 |

**TABLE II:** Table showing the selected time-course where higher activation occurred for ToM and pain networks.

| ToM Clips Description |  | Pain Clips Description |  |
| --- | --- | --- | --- |
| Sr. No. | Description | Sr. No. | Description |
| T1 | Peck flies away to happy cloud | P1 | Gus pulls porcupine spines from Peck's head |
| T2 | Peck caught gazing at happy clouds | P2 | Alligator biting Peck |
| T3 | Baby crying, then happy | P3 | Peck tossing porcupine |
| T4 | Peck dons gear to show why he left | P4 | Cloud makes animals (lightning) |
| T5 | Pan from happy clouds to lonely cloud (Gus) | P5 | Gus makes alligator (lightning) |

## II. Methods and Materials

### A. fMRI Movie watching Data set and MRI Preprocessing

In the current study, we analyzed the early childhood dataset, which contained 122 childhood samples (ranging from 3-12 yrs) and 33 adult samples (Total = 155) [5]. The data is available on the OpenfMRI database under the accession number ds000228. All the participants were from the local community and had submitted written consent from parent/guardian. The data were collected with approval from the Committee on the Use of Humans as Experimental Subjects (COUHES) at the Massachusetts Institute of Technology. Participants watched a sound-less short animated movie called “Partly Cloudy” for a total duration of 5.6 minutes, and the stimuli were validated to activate ToM and pain regions [24], [25], [5] (Refer Figure 2). 3-Tesla Siemens Tim Trio scanner was used to capture structural and functional images [26]. The dataset was preprocessed using SPM 8 and other toolboxes available for Matlab [27], which registered all functional images to the first run image and then registered that image to each participant’s structural images [5]. All structural images were normalized to Montreal Neurological Institute (MNI) template [28], [29]. The smoothening for all images was performed using a Gaussian filter and identified Artifactual timepoints using ART toolbox [30], [5].

**Figure. 1.**
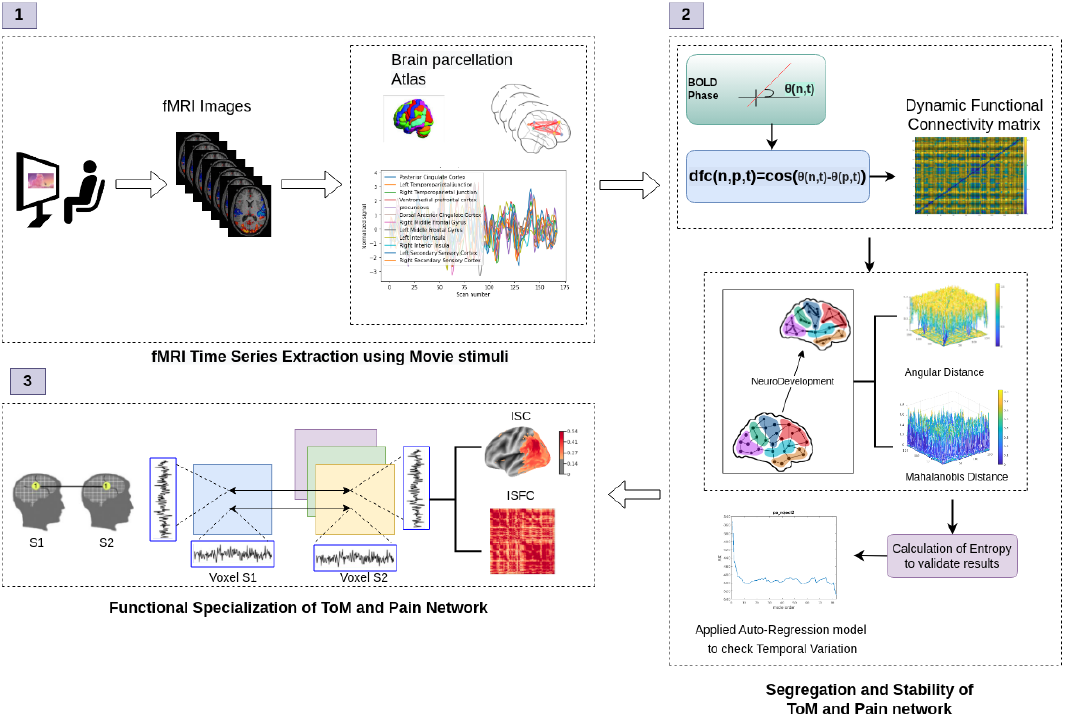
Illustrative overview of proposed architecture for identifying age at which ToM and pain networks are getting reasonable temporal stability and how it predicts behavioral scores of false-belief task. The paper’s contribution is as follows: **1)** Data collection and extraction of time-series from ToM and pain networks, **2)** Calculation of Angular distance and Mahalanobis distance using dominant dFC matrices, leading to temporal stability computations, **3)** Prediction of behavioral scores using ISC.

**Figure. 2.**
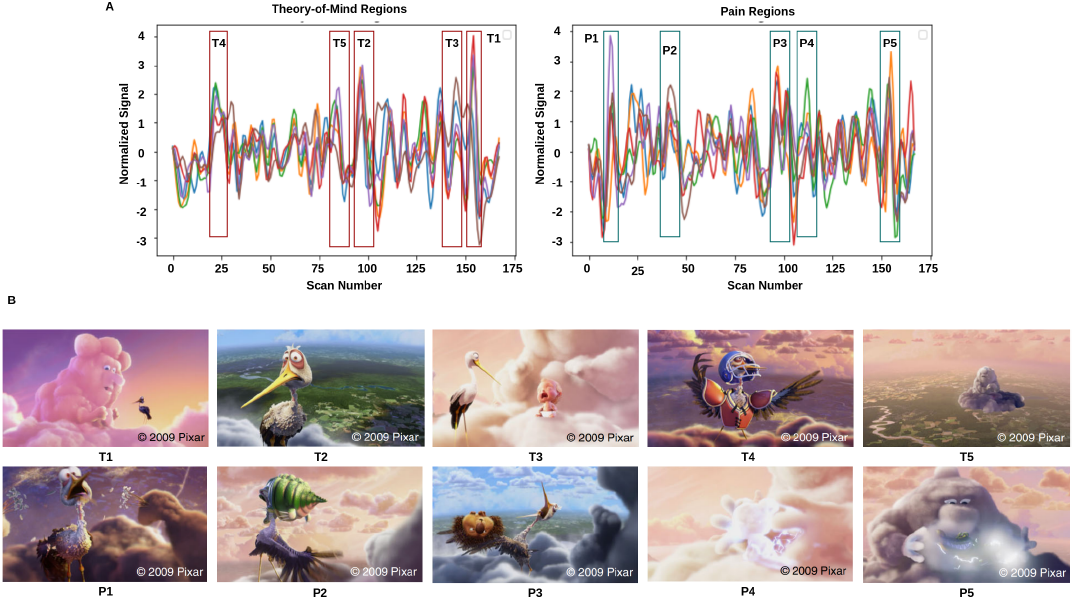
Movie demonstration: **A** represents the average time-series activation for the movie for all participants. **B** depicts the movie scenes where higher activation occurred. T1, T2, T3, T4, and T5 represent the ToM scenes with higher activation, whereas P1, P2, P3, P4, and P5 show the pain scenes with higher activation.

### B. Explicit ToM task and false-belief composite score and fMRI analysis

In the previous study, six-explicit ToM-related questions were administered for the false-belief task to identify the correlation between brain development and behavioral scores in ToM reasoning across a wide age range of children [4]. Each child’s performance on the ToM battery was assessed based on the proportion of correct answers from 24 matched items (14 prediction items and 10 explanation items). Based on the outcome of these explicit false-belief task scores, the participants were categorized into three classes: Pass (5-6 correct answers), inconsistent (3-4 correct answers), and fail (0-2 correct answers). In the current study, participant’s data were classified in two different ways: (a) into six groups of 3 years, 4 years, 5 years, 7 years, 8-12 years, and adults age groups (reference), and (b) into three false-belief tasks outcome-based groups, i.e., pass, inconsistent, and fail. The classification was undertaken to understand the differential developmental changes in the neural activation patterns [31]. Due to the unavailability of data on any 6-year-olds, that age group could not be added to the above categorization. Further, the BOLD time series for individual subjects from these groups were extracted for regions of interest (ROIs) anchored in two brain networks -Theory-of-Mind and Pain networks (Refer to Table 1). The regions selected for ToM were -bilateral Temporoparietal Junction (LTPJ and RTPJ), Posterior Cingulate Cortex (PCC), Ventro- and Dorso-medial Prefrontal Cortex (vmPFC and dmPFC), and Precuneus, whereas the Pain network regions consist of - bilateral Middle Frontal Gyrus (LMFG and RMFG), bilateral Interior Insula, and bilateral Secondary Sensory Cortex (LSSC and RSSC) [32]. These brain regions and their MNI coordinates were selected from published literature (Refer to Table 1) [33], [34]. We used the Schaefer atlas for brain parcellation, and by using MNI coordinates, we created a spherical binary mask with a 10 mm radius for all selected ROIs. Finally, Time series were extracted for each participant for the six regions of ToM, the six regions of Pain, and twelve brain regions of interest to test our hypothesis. Further analyses were carried out on the extracted time series signals.

### C. BOLD phase coherence and estimation of Dynamic Functional Connectivity

Functional connectivity (FC) is a widely used measure of brain connectivity that infers statistically significant co-activation patterns for a pair of brain regions. FC is estimated from Blood Oxygen Level-Dependent (BOLD) fMRI signals among pair of brain regions [35], [36], [37], [38]. However, these correlation and covariance measures assume that the time series signal for specific brain regions remains static over time, which significantly limits our understanding of whole brain dynamics associated with neurodevelopment. Further temporal changes could carry distinct and meaningful connectivity patterns over the scan and developmental time scales across brain regions [39], [40].

Resting-state data and complex naturalistic stimuli have been shown to encode significant variation among functional brain networks over the entire stimulus duration. Hence, dynamic functional connectivity (dFC) patterns across the whole data set can provide a more penetrating view into the activation patterns over the more commonly used static functional connectivity measure [41]. We have chosen an instantaneous measure for the computation of dFC to circumvent the temporal resolution issues that arise in the case of the more popularly employed sliding window correlation method, which is limited by the heuristic selection of the window size. Shorter windows include spurious correlations with high variability and low reliability and have a lower statistical significance due to a lesser number of data points, whereas longer windows are capable of eliminating noise-related correlation while failing to capture significant transient changes in the time series signals [42], [43], [41], [44]

Here, we employ BOLD signal Phase Coherence and quantify the strength of this synchronization while discarding its amplitude. The phase component of the signal sufficiently captures the temporally transient changes, which follows from the observation that two weakly-coupled nonlinear oscillators can synchronize even without any correlation of their amplitudes. Further, phase coherence does not assume stationarity of signals compared to other transformation and coherence-based methods, making this an ideal unsupervised method for characterizing dFC.

Finally, BOLD phase coherence was employed to estimate time-resolved dFC for each subject, resulting in a high dimensional N x N x T matrix (where N = 12 denotes the number of brain regions and T = 168 represents the total number of time points). The Hilbert transform was applied to the BOLD time series to compute the BOLD Phase Coherence to reduce dimensionality to enumerate the instantaneous phases *θ*(*n, t*) (Refer to Figure 2). The modulated BOLD signal s(t) was then represented analytically using the following equations [15]:

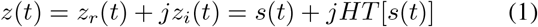

where *HT* [*∗*] stands for Hilbert Transformation. Using the following formulae, the instantaneous phase *θ*(*t*) was calculated [15].

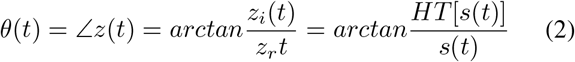

dFC (n,p,t) was then calculated for the predetermined brain areas n and p, as follows [15]:

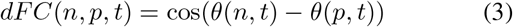

### D. Computation of Dominant Dynamic Functional Connectivity Subspaces

One major drawback is the inclusion of high-frequency noise, which can be resolved by narrow-band filtering or using an unsupervised dimensionality reduction method that excludes the noise and retains meaningful signals, such as the decomposition of the signal phase data using Principal Components Analysis (PCA). Applying PCA leading eigenvectors were estimated and arranged according to their percentage contribution to the total variance estimated from data [45].

A participant-specific dFC (n, p, t) matrix of dimension N X N X T reflecting the FC between the nth and pth brain region at each time point was subjected to principal component analysis (PCA). As a result, *dFC*(*n, p, t*) or simply *dFC*_*t*_ may be written as:

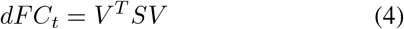

S stands for the diagonal matrix, and V stands for the leading eigenvector of the NXN matrix. The primary components of the dFC are represented by the number k=2. The dominating dFC D(n,k,t) was calculated as follows:

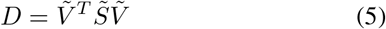

where 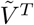 denotes the reduced matrix. Here, we choose k=2 or two leading eigenvectors that contain 80% information.

### E. Computation of Network Temporal Stability using Dominant dFC Subspaces

We characterize temporal stability dominating subspaces by estimating their similarity between different time points. Additionally, quantifying the temporal stability of a network throughout data acquisition can also vouch for the reliability of the connectivity observed and the robustness of the network activity in the face of external disturbances. We used two techniques to achieve that goal: Mahalanobis distance and Angular distance (Refer to Figure 2). We used the following equation to determine the distance between dFC sub-spaces from different time points in the principal angular distance method [15]:

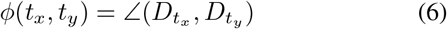

where each entry of the temporal stability matrix is *φ*(*t*_*x*_, *t*_*y*_), the range of angular distance is from 0 to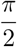, where 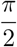,denotes high angular distance, and 0 denotes low angular distance. We calculated the angular separation between each individual’s *t*_*x*_ and *t*_*y*_ dominant dFC subspaces by calculating their angular separation, yielding a T X T matrix per subject. Finally, we estimated Euclidean distance using the Mahalanobis distance method, which used the following equation to measure the distance between distinct points from one space to another. It was done for each participant’s dFC dominating subspaces:

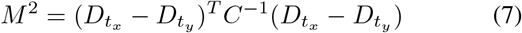

Where *M* ^2^ stood for the distance between each point, According to this method, the Euclidean distance in a 2D space is calculated between each ROI and the average of the rest of the ROIs between the dominant dFC subspaces at two different time points. [46]. We calculated a temporal stability matrix for each participant, where the Mahalanobis distance ranged from 0.5 to 2.5. Lower values indicate that subspaces were comparable, while larger values indicate that subspaces were distinct.

### F. Validation of Results

#### 1) Entropy

To validate the results of temporal stability matrices, we calculated entropy that defined measurement of detectable temporal order that we may interpret as the overall stability of the temporal stability matrices [15]. The lower the entropy value, the higher the instability in the temporal patterns and vice versa. For each subject, we compute the entropy of temporal stability matrices, where each element is the estimated Mahalanobis or Angular distance between the dominant subspaces *D*_*t*_*x* and *D*_*t*_*y* . The Entropy was calculated using the following formula [15]:

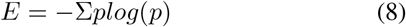

Where p holds the normalized histogram counts.

#### 2) Frobenius norm

The matrix’s Frobenius or Euclidean norm was utilized to quantify the variations between the temporal stability matrices generated for different age groups. We calculated Frobenius norm using the following formula [15]:

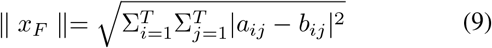

Where entries of temporal matrices are indicated by *a*_*ij*_ and *b*_*ij*_.

### G. Stochastic characterization of dFC

The degree of temporal changes in functional networks captured within the observed temporal fluctuations can be identified using the principal angle and the Mahalanobis distance between the dominant dFC subspaces [15]. We use auto-regressive (AR) models to uncover the underlying stochastic properties of these measurements (Refer to Figure 1). *AR*(*ρ*) concept was implemented using following equation [15]:

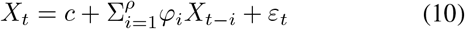

Where φ_*1*_*…* φ ρ=parameters of model, c= constant, *ε*_*t*_ = white noise, *ρ* = model order [15]. The Akaike information criterion may be used to calculate the ideal model of an AR process (AIC) using following equation [15]:

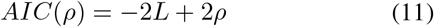

Where L is the likelihood function, we estimated several values for the model order parameter *AR*(*ρ*), ranging from 0 to 100, and then chose the AR model with the first minimum AIC score [47], [15], [48]. If it is determined that the model order is larger than 1, the underlying process is regarded as being non-Markovian.

### H. Quantifying Functional Network Specialization

To predict the behavioral score of participants for the false-belief task, we implemented Inter Subject Correlation (ISC), which measured responses activated during stimulus to naturalistic stimuli by taking into account solely the brain activity shared across all the subjects for the same stimulus [49]. The reason for using ISC was that when the participants were watching the movie, some brain areas were synchronized due to the same stimulus, and some subject-specific measurements contained idiosyncratic and non-stimulus-specific signals and noise [21], [22]. For example, the presence of substantial ISC in a particular region didn’t prove the stimulus activates that region; instead, it indicated that the region encodes information about the consistent stimulus across all individuals [22]. Significant ISC suggested that a region encodes information about a consistent stimulus across tasks or groups. If there is a strong inter-subject correlation (ISC), the reaction time course in one subject’s brain may predict that in another subject’s brain [22]. In this work, we computed leave-one-out ISC that calculated shared stimulus-related measures.

## III. Experimental Results

### A. Distinct Temporal Stability of ToM and Pain Network in 3-12 years age

#### 1) Computation of Temporal Stability using Angular Distance

To check the distinct temporal stability of ToM and Pain networks in 3-12 years of age, we first calculated angular distance matrices among dominant dFC subspaces identified over all the time points. Subsequently, a *time∗ time* temporal stability matrix was derived, which was then averaged over all individuals. Each entry in the matrix represented the angle between the dominating dFC subspaces at *t*_*x*_ and *t*_*y*_. As described earlier, we calculated Angular distance into two parts: one for six age categories and one for the three false-belief-task performance groups (Figure 3). Figures 3 and 4 represent the subject-average temporal matrices for all 12 ROIs and 6 ROIs in each ToM and Pain network. The higher the value of angular distance, the bigger the leap between two dominant dFC subspaces at time points *t*_*x*_ and *t*_*y*_. In Figure 3, we see higher high-distance data points in the matrices 3 yrs and 4 yrs, which subsequently reduces and stabilizes for adults (reference group). The 8-12 years matrix shows high distance values due to the inclusion of both ToM and Pain network ROIs and a wide range of ages, a limitation peculiar to the acquired dataset. In Figure 4, the pattern of high distance values indicates lower overall distance values as we progress through the ages. The only anomalies in this trend seen in the three figures are the slight increase in switching activity from 5 yrs to 7 yrs and an increase in distance values from 3 yrs to 4 yrs. The second parameter for qualitative interpretation is the stability of the temporal distances, i.e., how long a certain distance value persists during the observation period. The persistence of distance value patterns alludes to the stability of a system. In particular, 3yrs and 4yrs children’s data exhibits frequent temporal switching activity with very low temporal stability. The temporal matrices per age group show higher distance values for All ROIs than the corresponding matrices for ToM or Pain ROIs, and this difference persists through age. We also found more stability in ToM scene activation for 3 yrs, 4 yrs, 5 yrs, and 7 yrs age groups. For stability in pain scene activation, 3 yrs, 5 yrs, 7 yrs, and 8-12 yrs age groups showed higher temporal stability (see Figs 2, 4).

**Figure. 3.**
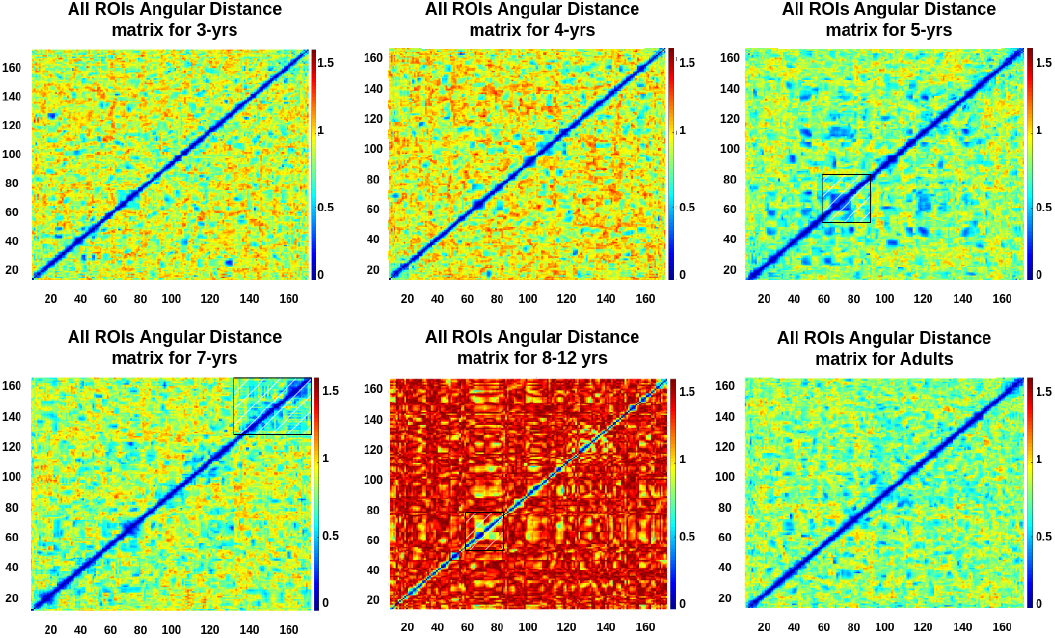
Angular distance matrices depicting temporal patterns of all ROIs of the ToM and Pain networks for six age groups (3, 4, 5, 7, 8-12-year children and Adults). The figure shows less temporal stability at 3 yrs and 4 yrs, and considerable temporal stability at 5 yrs, and so on. The higher angular distance for the 8-12 yrs age group is due to the inclusion of samples from discontinuous age groups

**Figure. 4.**
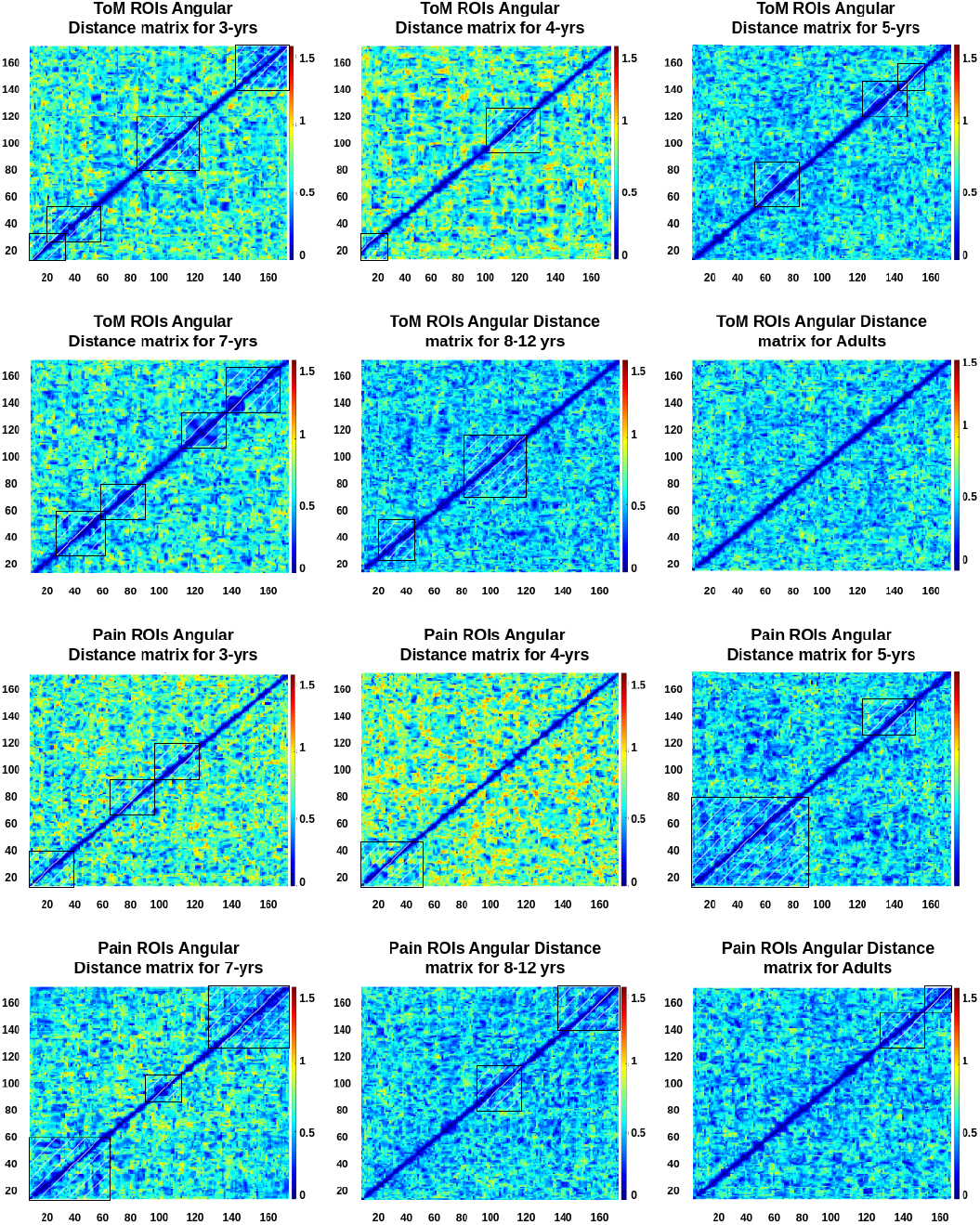
Angular distance matrices depicting temporal patterns of ToM and pain network ROIs for each of the six age groups (3, 4, 5, 7, 8-12 yrs and Adults age groups). We observed temporal stability in ToM and pain networks at age 3 yrs for ToM and pain movie scene activation.

To quantify differences between temporal matrices, the distance values were converted into Z-scores, and Kolmogorov-Smirnov tests for equality of distributions (all groups were found to have unequal distributions with *p* varying from < 0.4 to < 4 ×10^−313^), followed by Kruskal Wallis tests were conducted: All ROIs *χ*^2^(5) = 42.04, *p* < 6 ×10^−8^; Pain ROIs *χ*^2^(5) = 47.63, *p* < 5× 10^−9^; ToM ROIs *χ*^2^(5) = 24.59, *p* < 0.0003. Dunn-Sidak post-hoc test was performed for pairwise comparisons: All ROIs: 3yrs-4yrs: diff=-1376.4, *p* = 0.0114, 4yrs-5yrs: diff=1716.9, *p* = 4.8131 ×10^−4^, 4yrs-8yrs: diff=2212.4, *p* = 1.3683 ×10^−6^, 4yrs-Adult: diff=2004.2, *p* = 1.9045 ×10^−5^, 7yrs-8-12yrs: diff=1551.7, *p* = 0.0025, 7yrs-Adult: diff= 1343.6, *p* = 0.0149; Pain ROIs: 3yrs-8-12yrs: diff= 1365.7, *p* = 0.0125, 3yrs-Adult: diff=1196.6, *p* = 0.0445, 4yrs-5yrs: diff=1735.3, *p* = 3.9715 ×10^−4^, 4yrs-8-12yrs: diff=2337.3, *p* = 2.6373× 10^−7^, 4yrs-Adult: diff=2168.2, *p* = 2.4350× 10^−6^, 7yrs-8-12yrs: diff=1556.7, *p* = 0.0023, 7yrs-Adult: diff=1387.6, *p* = 0.0104; ToM ROIs: 3yrs-8-12yrs: diff=1286.6, *p* = 0.0232, 4yrs-8-12yrs: diff=1836.4, *p* = 1.3342 ×10^−4^, 4yrs-Adult: diff=1319.2, *p* = 0.0180, 7yrs-8-12yrs: diff=1302.9, *p* = 0.0205; rest NS. To quantify the complexity of these temporal stability patterns, entropy was calculated. Figure 5 shows the entropy values calculated from angular distance values for the six age groups as violin plots. In all three graphs (Fig 5 subgraphs -A, E, F), a higher entropy value is noticed for 4 yrs compared to the other age groups. A lower entropy value is noticed for 7 yrs group as compared to the 5 yrs group, and this corroborates with the qualitative conclusions drawn from the temporal matrices earlier. Kolmogorov-Smirnov test was performed to check for equality of distributions (all groups were found to have unequal distributions with *p* < 0.05), followed by the Kruskal-Wallis tests with non-significant results at 5% confidence level (All ROIs *χ*^2^(5) = 5.54, *p* < 0.4; Pain ROIs *χ*^2^(5) = 10.97, *p* < 0.06; ToM ROIs *χ*^2^(5) = 4.13, *p* < 0.6).

**Figure. 5.**
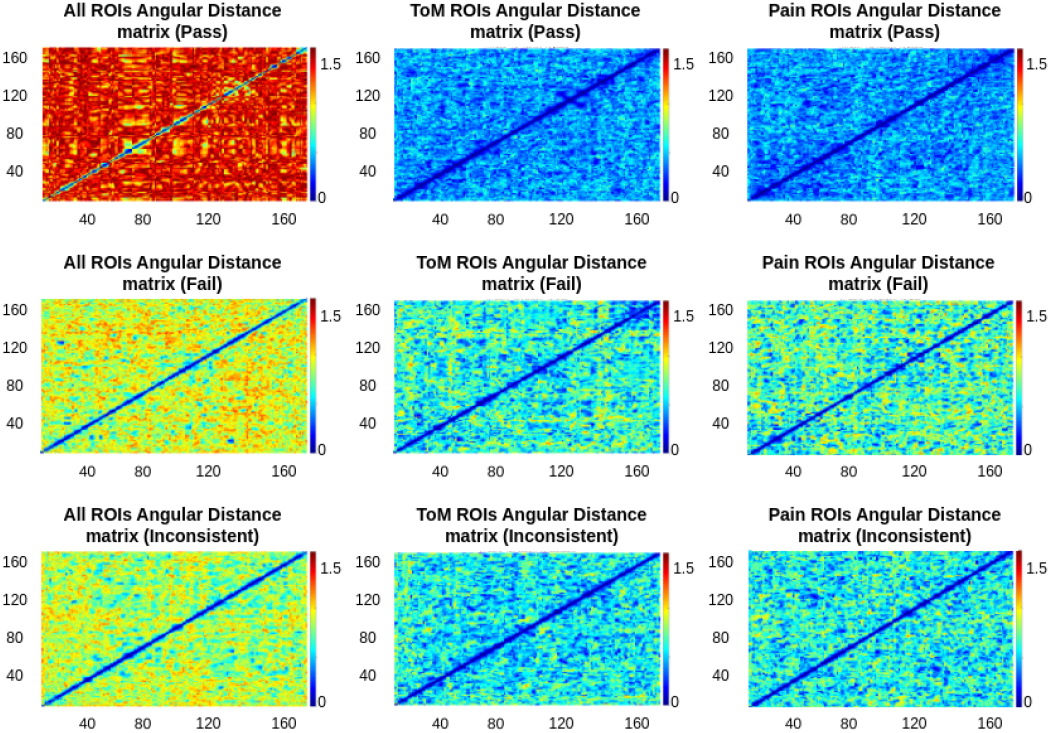
Angular distance matrices depicting temporal patterns for false-belief task-based pass, fail, and inconsistent groups. It showed false-belief task is dependent on temporal stability. The figure shows higher temporal stability in ToM network for the pass group, moderate stability for the inconsistent group, and lowest temporal stability for the fail group.

**Figure. 6.**
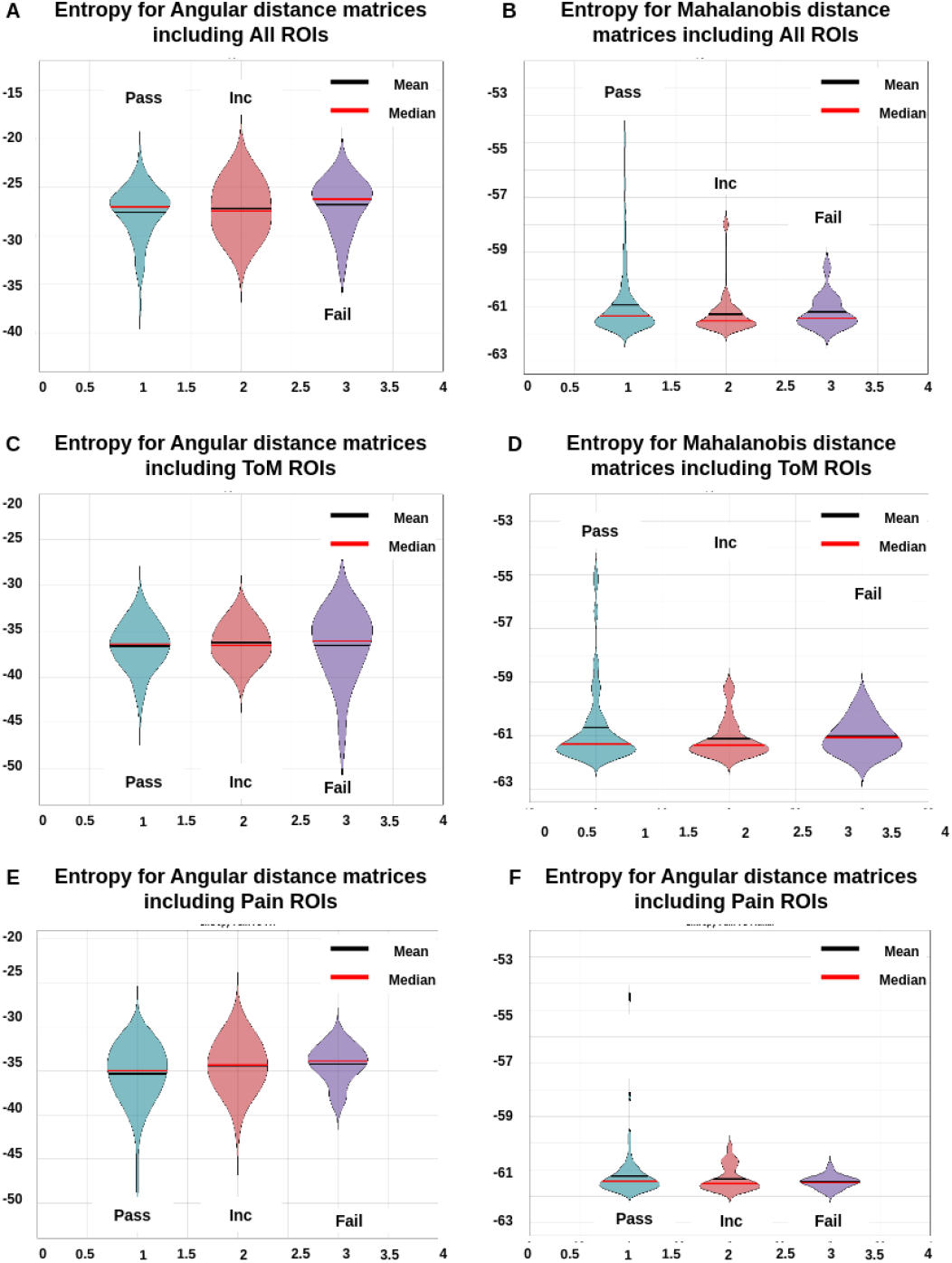
**A, C, E** depicts entropy plots for angular distance for the false-belief task-based pass, fail, and inconsistent groups. **B, D, F** are depicting entropy plots for Mahalanobis distance. In the case of entropy plots, Mahalanobis distance is better at capturing the differences in temporal stability in different groups.

In addition, we computed the Frobenius distance to examine the difference among temporal matrices of ToM and Pain regions (Figure 4). We find Frobenius distance values to progressively reduce from 3 yrs to Adults, with an anomalous dip for 5 yrs. No statistical significance was found using the Kruskal-Wallis test (*χ*^2^(5) = 7.59, *p* < 0.2).

#### 2) Computation of Temporal Stability using Mahalanobis

*Distance:* Next, we estimated the Mahalanobis distance to assess the temporal stability of the dFC. Each matrix element represents the Mahalanobis distance between the dominating dFC subspaces (Refer to Supplementary Material Section 1). A higher value of Mahalanobis Distance denotes a larger distance between the average of the Euclidean distance between individual ROIs at *t*_*x*_ and the collection of all ROIs (ToM or Pain network ROIs) at *t*_*y*_ of the dominant dFC subspaces. The above can be interpreted similarly to the Angular Distance measure. We observe a trend of decreasing high distance values to lower overall distance values from children to adults. Consistent with the previous measure, we also find a slight increase in temporal switching dynamics from 5 yrs to 7 yrs and an increase in distance values from 3 yrs to 4 yrs. A major reason for the limited inference drawn from these graphs is the presence of outlying data points with high distance values, which obscures smaller distance values.

Z-scores were calculated to compare the temporal matrices of these subgroups, Kolmogorov-Smirnov test was performed (all groups were found to have unequal distributions with *p* varying from < 2 ×10^−8^*to* < 1 ×10^−255^), followed by Kruskal-Wallis (All ROIs *χ*^2^(5) = 18066.11, *p* = 0; Pain ROIs *χ*^2^(5) = 3863.9, *p* = 0; ToM ROIs *χ*^2^(5) = 5254.75, *p* = 0) and Dunn-Sidak tests (Refer to Table 4).

To validate the results, we calculated entropy (shown in Figure 5), with similar entropy values seen across the age groups, with an exception for ToM ROIs, which displays a pattern similar to the observations from the angular distance measure: a higher entropy (and hence more complexity) in4yrs as compared to 3 yrs, and in 7 yrs as compared to 5yrs. Kolmogorov-Smirnov test was performed to check for equality of distributions (all groups were found to have unequal distributions with *p* < 0.05), followed by the Kruskal Wallis tests (All ROIs *χ*^2^(5) = 72.57, *p* < 3 ×10^−14^; Pain ROIs *χ*^2^(5) = 77.87, *p* < 2.5 × 10^−15^; ToM ROIs *χ*^2^(5) = 82.44, *p* < 3× 10^−16^) and Dunn-Sidak for the significant results of the six age subgroups (All ROIs: 3yrs-Adult: diff=-82.5009, *p* = 3.1472 × 10^−8^, 4yrs-Adult: diff=-72.7089, *p* = 5.6670 × 10^−6^, 5yrs-Adult: diff=-77.3391, *p* = 2.0698 × 10^−8^, 7yrs-Adult: diff=-76.595, *p* = 2.5474 × 10^−8^, 8-12yrs-Adult: diff=-64.2068, *p* = 9.1637 × 10^−8^; Pain ROIs: 3yrs-Adult: diff=-77.9412, *p* = 1.0955 × 10^−7^, 4yrs-Adult: diff=-76.0714, *p* = 1.6206 × 10^−6^, 5yrs-Adult: diff=-81.3824, *p* = 2.0677 × 10^−8^, 7yrs-Adult: diff=-77.0435, *p* = 2.4449×10−8, 8-12yrs-adult: diff=-74.2941, *p* = 2.0845 × 10^−8^; ToM ROIs: 3yrs-Adult: diff=-77.1301, *p* = 1.4829 × 10^−7^, 4yrs-Adult: diff=-75.5671, *p* = 1.9607 × 10^−6^, 5yrs-Adult: diff=-83.0419, *p* = 2.0676 × 10^−8^, 7yrs-Adult: diff=-58.8590, *p* = 2.0455 × 10^−5^, 8-12yrs-Adult: diff=064.6889, *p* = 7.5010 ×10^−8^; rest NS).

We also calculated the Frobenius distance between the ToM and Pain network ROIs for the six age subgroups (see Fig 5). We observe a higher distance for 3 yrs and Adults and a lower distance for all the other age groups. No statistical significance was found using the Kruskal-Wallis test (*χ*^2^(5) = 7.78, *p* < 0.2).

### B. Stochastic Characterization of Temporal Stability Measures

Next, we investigated the stochastic characteristics of dFC development by using the principal angle *φ*(*t*) and Mahalanobis distance M(t) as time functions. The temporal changes in *φ*(*t*) and M(t) are defined as autoregressive processes to yield the function: *AR*(*ρ*). Further, the first minimum AIC score for the generated AR models is considered the best-fit model order. We found the optimum model order of *ρ≥* 2 for Angular distance and *ρ ≥*4 for Mahalanobis distance (Refer to supplementary material Table 9 and Fig 5).

### C. Predicting False-belief Task-Based Pass, Fail, and Inconsistent Groups based on ISC and Temporal stability analysis

Next, we tested our hypothesis that the temporal stability patterns of social brain regions of ToM in 3-12 years could predict behavioral scores of the False-belief task. Temporal instability was a dominant feature at 3 and 4 yrs, but at age 5, we discovered higher temporal stability (Refer to Figs 3 and 4). In the current dataset, out of 122 participants (age ranges 3-12 yrs), 15 participants failed the false-belief task. They belonged to 3 yrs and 4 yrs age groups, whereas 23 participants were inconsistent during tasks and belonged to mostly 3 yrs and 4 yrs and few participants from 5 yrs. 84 participants passed the task; approximately all participants passed the task from 5 yrs age group onward. To validate our results to get more nuanced insight, we also performed temporal stability and Inter Subject Correlation (ISC) analysis for false-belief task-performers. We compared the results of these two measures. Our hypothesis accurately predicted participants who failed, passed and gave inconsistent responses in the False-belief task. For the participants who passed ToM task, their ISC scores were high for ToM network with low angular distance. In contrast, participants who failed the task’ ISC values were low for ToM network with high angular distance (Refer to Figures 4 and 9) (Refer to supplementary material section 2).

#### 1) Temporal Stability using Angular Distance

A qualitative analysis of the angular distance matrices for the three false-belief task performances (Figure 10) reveals a lower switching in dFC states and higher temporal stability for passers when all and only ToM and Pain ROIs are considered. This pattern destabilizes with higher distance values are seen for inconsistent performers, and the fail group has the highest instability (Refer to Figures 7 and 8).

**Figure. 7.**
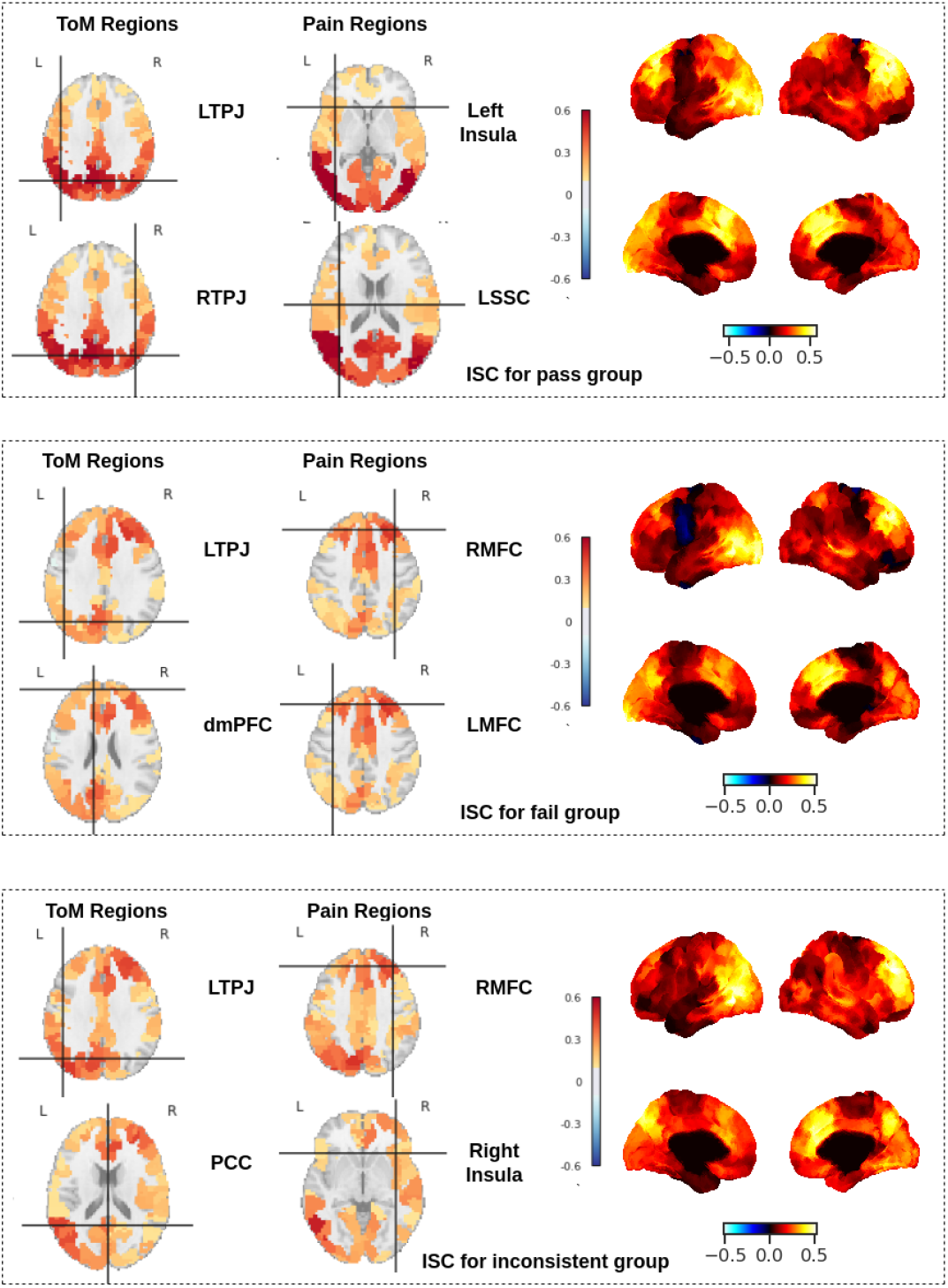
Mean ISC for a false-belief task-based pass, fail, inconsistent groups. It showed Neurodevelopment was dependent on developmental psychology. We found that in the group who passed the false-belief task, their ISC was high in ToM regions, whereas we observed moderate ISC for ToM regions in the inconsistent group and hypo ISC for the fail group.

To quantify differences between temporal matrices, the distance values were converted into Z-scores, and Kolmogorov-Smirnov tests for equality of distributions (all groups were found to have unequal distributions with *p* varying from < 0.4 to < 4 ×10^−313^), followed by Kruskal Wallis tests were conducted: All ROIs *χ*^2^(2) = 7.31, *p* < 0.03; Pain ROIs *χ*^2^(2) = 15.63, *p* < 0.0005; ToM ROIs *χ*^2^(2) = 12.18, *p* < 0.003. Dunn-Sidak posthoc test was performed for pairwise comparisons: All ROIs: Pass-Fail: diff=-533.0409, *p* = 0.0271; Pain ROIs: Pass-Inc: diff=-524.7799, *p* = 0.0302, Pass-Fail: diff=-806.1150, *p* = 2.8977 ×10^−4^; ToM ROIs: Pass-Fail: diff=-722.1818, *p* = 0.0014; rest NS.

To quantify the complexity of these temporal stability patterns, entropy was calculated. Kolmogorov-Smirnov test was performed to check for equality of distributions (all groups were found to have unequal distributions with *p* < 0.05), followed by the Kruskal Wallis tests with non-significant results at 5% confidence level (All ROIs *χ*^2^(2) = 1.38, *p* < 0.6; Pain ROIs *χ*^2^(2) = 2.11, *p* < 0.4; ToM ROIs *χ*^2^(2) = 2.11, *p* < 0.8).

#### 2) Temporal Stability using Mahalanobis Distance

Qualitative analysis of the Mahalanobis distance matrices reveals no differences for the False-Belief performances (Refer to Supplementary Materials Figure 8)

We calculated Z-scores for comparing the temporal matrices of pass, fail, and inconsistent groups; the Kolmogorov-Smirnov test was performed, followed by Kruskal-Wallis (All ROIs *χ*^2^(2) = 3171.04, p = 0; Pain ROIs *χ*^2^(2) = 5558.54, p = 0; ToM ROIs *χ*^2^(2) = 4777.35, p = 0) and Dunn-Sidak tests (Refer to Table 3). We also calculated entropy and found an increasing trend for the false-belief test performers, denoting increasing instability from passers to inconsistent performers and failers. To check equality of distribution, we performed the Kolmogorov-Smirnov test and found all false-belief task groups with unequal distribution with *p* < 0.05 (All ROIs *χ*^2^(2) = 0.61, *p* < 0.75; Pain ROIs *χ*^2^(2) = 0.57, *p* < 0.8; ToM ROIs *χ*^2^(2) = 0.71, *p* < 0.75).

**TABLE III:** Table with difference and p-values calculated using the posthoc pair-wise Dunn-Sidak test for Mahalanobis Distance-derived Z-scores for the six age subgroups (3, 4, 5, 7, 8-12 yrs and Adult), and for the False-Belief Performance results.

| Pairs | All ROIs |  | Pain ROIs |  | ToM ROIs |  |
| --- | --- | --- | --- | --- | --- | --- |
|  | Diff. | p-value | Difference | p-value | Difference | p-value |
| 3yrs-4yrs | 53917 | $2.0676 \times 10^{-8}$ | 4719.4 | $2.0676 \times 10^{-8}$ | 23518 | $2.0676 \times 10^{-8}$ |
| 3yrs-5yrs | 31162 | $2.0676 \times 10^{-8}$ | -16415 | $2.0676 \times 10^{-8}$ | -1104.3 | 0.0818 |
| 3yrs-7yrs | 18966 | $2.0676 \times 10^{-8}$ | -8762 | $2.0676 \times 10^{-8}$ | 8240.2 | $2.0676 \times 10^{-8}$ |
| 3yrs-8-12yrs | 24470 | $2.0676 \times 10^{-8}$ | -9564.9 | $2.0676 \times 10^{-8}$ | -1192.5 | 0.0458 |
| 3yrs-Adult | 29884 | $2.0676 \times 10^{-8}$ | -13855 | $2.0676 \times 10^{-8}$ | 6363.1 | $2.0676 \times 10^{-8}$ |
| 4yrs-5yrs | -22755 | $2.0676 \times 10^{-8}$ | -21135 | $2.0676 \times 10^{-8}$ | -24623 | $2.0676 \times 10^{-8}$ |
| 4yrs-7yrs | -34951 | $2.0676 \times 10^{-8}$ | -13481 | $2.0676 \times 10^{-8}$ | -15278 | $2.0676 \times 10^{-8}$ |
| 4yrs-8-12yrs | -29447 | $2.0676 \times 10^{-8}$ | -14284 | $2.0676 \times 10^{-8}$ | -24711 | $2.0676 \times 10^{-8}$ |
| 4yrs-Adult | -24033 | $2.0676 \times 10^{-8}$ | -18574 | $2.0676 \times 10^{-8}$ | -17155 | $2.0676 \times 10^{-8}$ |
| 5yrs-7yrs | -12197 | $2.0676 \times 10^{-8}$ | 7653.3 | $2.0676 \times 10^{-8}$ | 9344.4 | $2.0676 \times 10^{-8}$ |
| 5yrs-8-12yrs | -6692 | $2.0676 \times 10^{-8}$ | 6850.4 | $2.0676 \times 10^{-8}$ | -88.2839 | 0.9999 |
| 5yrs-Adult | -1278 | 0.0247 | 2560.7 | $2.9612 \times 10^{-8}$ | 7467.4 | $2.0676 \times 10^{-8}$ |
| 7yrs-8-12yrs | 5504.7 | $2.0676 \times 10^{-8}$ | -802.8698 | 0.3779 | -9432.7 | $2.0676 \times 10^{-8}$ |
| 7yrs-Adult | 10919 | $2.0676 \times 10^{-8}$ | -5092.6 | $2.0676 \times 10^{-8}$ | -1877 | $8.4552 \times 10^{-5}$ |
| 8-12yrs-Adult | 5414.1 | $2.0676 \times 10^{-8}$ | -4289.7 | $2.0676 \times 10^{-8}$ | 7555.7 | $2.0676 \times 10^{-8}$ |
| Pass-Inc | 11114 | $9.5606 \times 10^{-10}$ | 7003.8 | $9.5606 \times 10^{-10}$ | 10777 | $9.5606 \times 10^{-10}$ |
| Pass-Fail | 2515.5 | $9.5606 \times 10^{-10}$ | 15410 | $9.5606 \times 10^{-10}$ | 13537 | $9.5606 \times 10^{-10}$ |
| Inc-Fail | -8598.4 | $9.5606 \times 10^{-10}$ | 8406.7 | $9.5606 \times 10^{-10}$ | 2759.2 | $9.5606 \times 10^{-10}$ |

#### 3) Functional Brain Network Specialization for False-Belief Task-Based Groups

For false-belief task-based pass, fail, and inconsistent groups, we found that in the participants who passed the task, their ToM regions were highly activated (i.e., PCC (0.6, p-value *<*0.02), LTPJ (0.6, p-value *<*0.01)), RTPJ (0.5, p-value < 0.025)), Precuneus (0.4, p-value *<*0.04)) and pain regions were moderately connected (average pain ISC= 0.17 (p-value < 0.045)). The participants who failed false-belief tasks in their ToM regions were not activated ((average ToM ISC= 0.12 (p-value < 0.05), average pain ISC= 0.16 (p-value < 0.04)). In contrast, who were inconsistent during false-belief tasks, their ToM regions were moderately connected ((average ToM ISC= 0.28 (p-value < 0.02), average pain ISC= 0.2 (p-value < 0.025)) (Refer to Figure 9).

## IV. DISCUSSION

Children can understand others’ desires, thoughts, and emotions during early brain development and distinguish bodily pains and reflexes. For example, what broadly encompasses social cognition, young infants acquire a remarkably sophisticated understanding of others’ intentions, ideas, and emotions, as opposed to their physiological effects, sensations, and diseases; most of this development happens before children begin conventional education at the age of six [50], [51], [52], [4]. Although adult, adolescent, and older children’s brain areas implicated in ToM have been widely investigated, fMRI investigations provide significant challenges for early childhood data. One of the key objectives of this study was to use a naturalistic task that included significant ToM and pain network activations during movie watching to track their temporal brain dynamics during early development. Additionally, to identify at what age reasonable temporal stability was manifested and how the development of ToM ability could be predicted over time. We estimate the temporal dynamics of dFC matrix in ToM and Pain networks and the separation between dFC subspaces based on angular distance and Mahalanobis distance measure to quantify the temporal stability of these cortical networks. Next, we test the hypothesis that the temporal stability and complexity estimated by Entropy are key measures that methodologically track brain maturation in children. In particular, the emergence of children’s mental reasoning and concept-understanding ability with increasing age. Their ability to reason about cartoon character’s bodies (the pain sensations) versus minds (the theory of mind network) is strongly associated with the acquired temporal stability of ToM and Pain functional brain networks. Hence, our critical finding bridges the gap between temporal dynamics estimated using frequently reported neuroimaging studies in older children and several important behavioral studies capturing the development of ToM reasoning ability.

We hypothesized an association between temporal stability and behavioral scores, i.e., temporal stability could predict the performance of false-belief task-based groups. We included a dataset in which participants aged 3 to 12 and adults watched a short animated movie (naturalistic stimuli) [4]. For example, one of the movie clips consists of Pixar’s ‘Partly Cloudy’ (see Figure 2), depicting multiple events capturing two major aspects (bodily sensations (often physical pain) and their mental states (beliefs, desires, and emotions)) of the main characters (a cloud named Gus and his stork friend Peck) activating ToM and pain sensory networks [4]. The dataset was first segmented into 3 yrs, 4 yrs, 5 yrs, 7 yrs, 8-12 yrs, and adults to cover continuous changes and then into three false-belief task-based pass, fail, and inconsistent groups to check the convergence of findings between predictions from brain development and what is generally known from experimental psychology.

In a previous study, the authors investigated the development of ToM and pain network using the static functional connectivity approach focusing on static correlation patterns between regions[4]. However, as participants watched the movie stimulus with engaging narratives, therefore, not only did brain networks show particularly dramatic change with age but also dynamic stability during stimulus-induced activation patterns; hence, a dynamic perspective is much warranted to get additional fundamental insights about the complex developmental changes across ages[4], [53], [54], [15]. Our main contribution to this article is applying dynamic techniques to demonstrate that temporal stability shows dramatic change with age in children and carries distinctive signatures for ToM and Pain networks. ToM stability was captured by decreasing angular distance and increasing stability from 3 yrs old children to adults, with 4 yrs and 7 yrs showing higher angular distance and instability and 5 yrs showing a pattern similar to that of the 8-12 yrs age group. The higher stability of higher-cognition and associative brain regions (included in the ToM and Pain networks) are useful in conferring adaptability and increased capacity to coordinate information processing across the cortical networks [55], [13], [53], [56]. Hence, our results suggest a higher stability, adaptability, and by extension, a higher propensity for mentalization of concepts by 5 yrs of age. This trend could not be observed as clearly in the Mahalanobis Distance analysis due to the smaller distance values caused by the high-value data outliers. The temporal matrices per age group show higher distance values for All ROIs than the corresponding matrices for ToM or Pain ROIs, and this difference persists through age.

For example, developing temporal differences between ToM and pain networks based on angular distance and Mahalanobis Distance analysis may reflect intrinsic developmental changes in brain networks and the emergence of functional selectivity of the event-driven response in individual brain regions. These findings may index distinctive features of stability of ToM and Pain networks from as early as 3 yrs of age. For the false-belief performance groups, a lower distance and higher repeatability in patterns were observed for passers, with increasing distances and instability observed for failers. From Frobenius Distance analysis, we find the lowest distance between ToM and Pain network activation patterns for 7 yrs, followed by slightly higher values for 5 yrs, 8-12 yrs, and Adults, high value for 3 yrs, and the highest value for 4 yrs. The literature found that ToM network is segregated from other networks at the early childhood stage and gets more functionally specialized throughout developmental time scales [4], [57], [56]. Future studies could provide more critical insight into the temporal stability of stimulus-induced versus task-driven networks to provide critical insights about spontaneous temporal stability patterns. However, collating such data on 3-year-old children will always remain challenging.

The second key hypothesis was to check the alteration between the temporal stability of ToM networks related to children’s ToM reasoning abilities [58], [59]. In the current dataset, the participants answered six questions about predicting and explaining actions based on false beliefs. The participants who failed in this task belonged to 3yrs and 4 yrs groups; interestingly, the same age groups exhibited low temporal stability (Refer to Fig 3), suggesting the new development of the concept around age four years. Moreover, participants who passed the test were mostly belonging to 5 yrs age group, where we observed reasonable temporal stability suggesting the development of transition from failure to success on the false-belief task is associated with the temporal stability of ToM brain networks. To obtain more clarity, we performed temporal stability analysis for pass, inconsistent, and fail groups separately and observed higher temporal stability for the pass group, moderate temporal stability for the inconsistent group, and low temporal stability for the fail group, These results were also validated by using multiple analysis including results based on entropy and other measures (Refer to Figs 7 and 8). Existing literature demonstrates that during the naturalistic movie-watching task, ISC measure predicts behavioral performance accurately [21], [22], [54]. Hence, we performed ISC analysis to predict false-belief task-based groups and their performance in the ToM task. We observed hyper ISC in the ToM network for the pass group, moderate ISC patterns for the inconsistent group, and hypo ISC for the failed group. We compared our findings from the temporal stability analysis. We found qualitative overlap with ISC results suggesting multiple analysis approaches could delineate a fundamental aspect of developmental differentiation in the social brain areas to depict others’ bodies versus minds. In summary, the current study provides a novel approach to predicting developmental stability and distinction in ToM versus Pain networks in the brain and accurate predictions based on brain features of behavioral performance in children. The distinct patterns of temporal stability of the two social cognition networks strongly predict the maturity of each network in response to the movie. Specific peak events within the movie evoke temporal dynamics, stability, and complexity that increases with age and the evolving theory of mind reasoning ability.

## V. CONCLUSION

In this research, we examined the developmental changes in the temporal stability of ToM and pain networks using a dynamic functional connectivity approach from childhood to adolescence. The study’s outcomes suggested that the ToM network acquires reasonable temporal stability in the early years, and similar dynamic patterns are observed for pain sensory networks. These dynamic stability patterns do not necessarily correspond with a loss of continuity in the neural basis at certain age independent of the major hallmark of ToM behavioral development, passing explicit false-belief tasks. Interestingly, particular time points during naturalistic movies where both ToM and pain networks exhibited higher stability. Functional brain networks are not well segregated from each other at the early childhood stage, and eventually, it gets more specialized with increasing age. This study has important limitations: 1) We used a dataset containing 155 subjects (33 adults) with naturalistic task data without resting state data, which should have been important for comparing spontaneous versus movie task-evoked stability in children. 2) In the future, we plan to continue investigating temporal dynamics with the availability of open datasets under different task conditions to test the generality of our findings. 3) Lastly, the association between two independent measures, temporal stability and ISC, needed to be sufficiently explored, as both were predictive of children’s behavioral performance, and their mathematical relationship remained an open question for future studies. Despite the above limitations, our current findings could be important for understanding atypical temporal stability in the impaired ToM ability in neurodevelopmental disorders (e.g., ASD, ADHD, etc.), thus opening new avenues for cognitive and developmental neuroscience.

## VI. APPENDIX

### A. Code Availability

Ex-AI model implementation will be made available on GitHub: https://github.com/dynamicdip/ The pipeline for the Analysis of Connectomes (C-PAC), including slice-time correction, motion correction, functional normalization, and smoothing procedure on ABIDE dataset including raw T1 and T2 images of fMRI resting state data are all available from http://preprocessed-connectomes-project.org/abide/.

## B. ACKNOWLEDGMENT

We acknowledge the generous support of IIT Jodhpur Core funds and the Computing facility. D.R. acknowledges the generous support of the NBRC Flagship program BT/MEDIII/NBRC/Flagship/Pro-gram/2019: Comparative mapping of common mental disorders (CMD) over the lifespan.

## Supplementary materials and methods

### ToM and Pain Network Segregation and Temporal Stability

#### Result for Six Age categories

**Figure 1:**
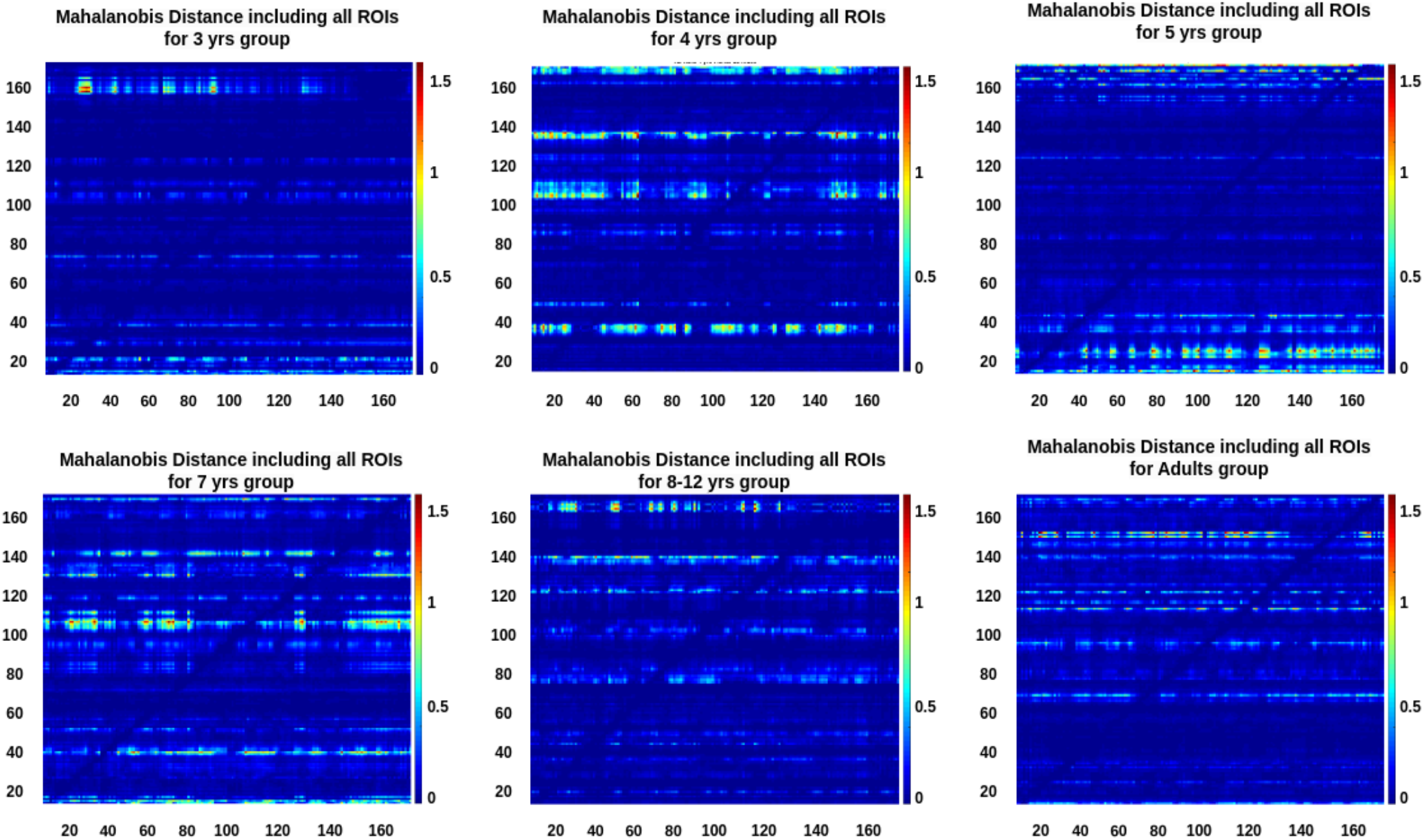
Mahalanobis distance for 6-groups including all ROIs

**Figure 2:**
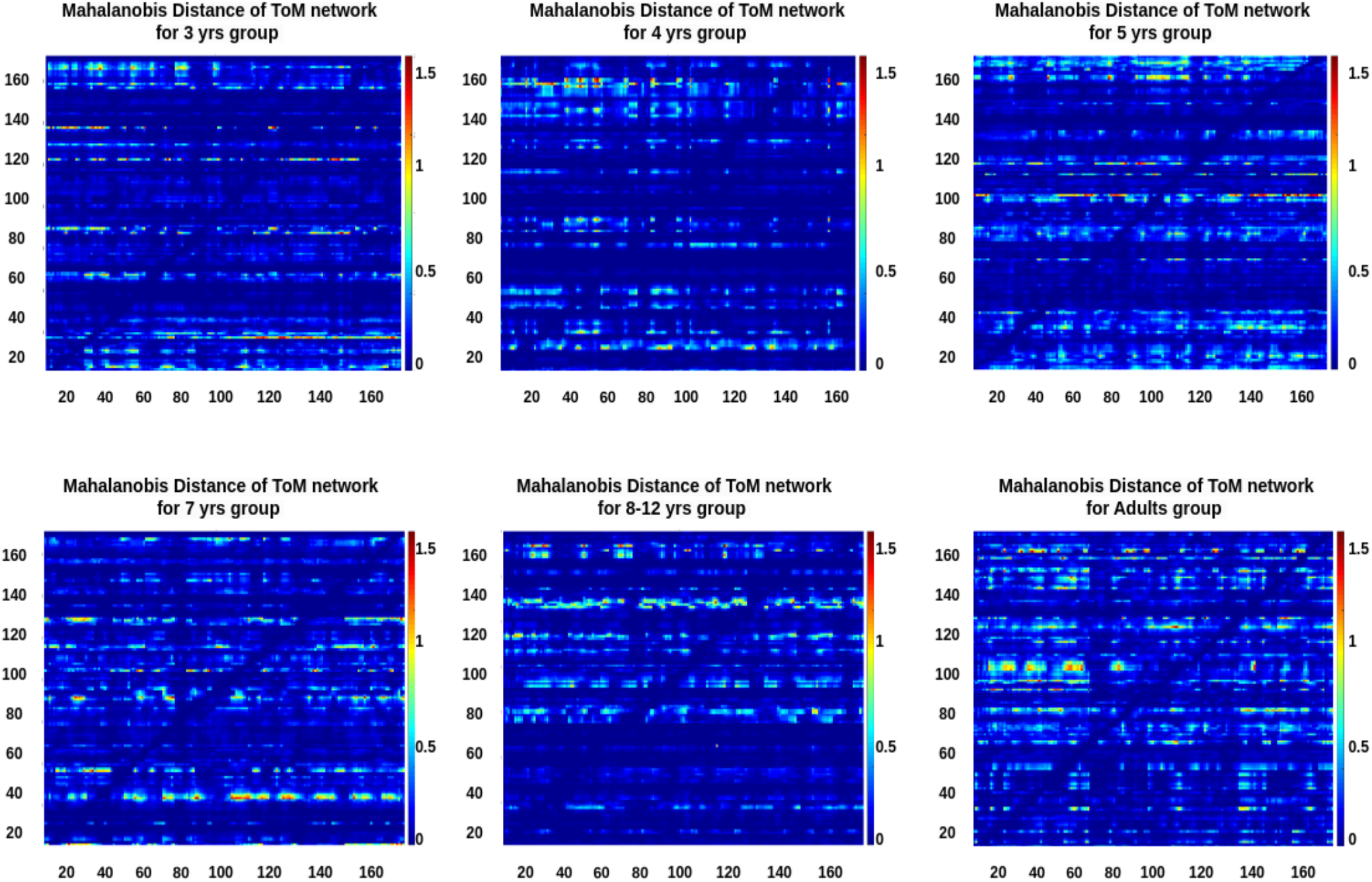
Mahalanobis distance for 6-groups including ToM ROIs

**Figure 3:**
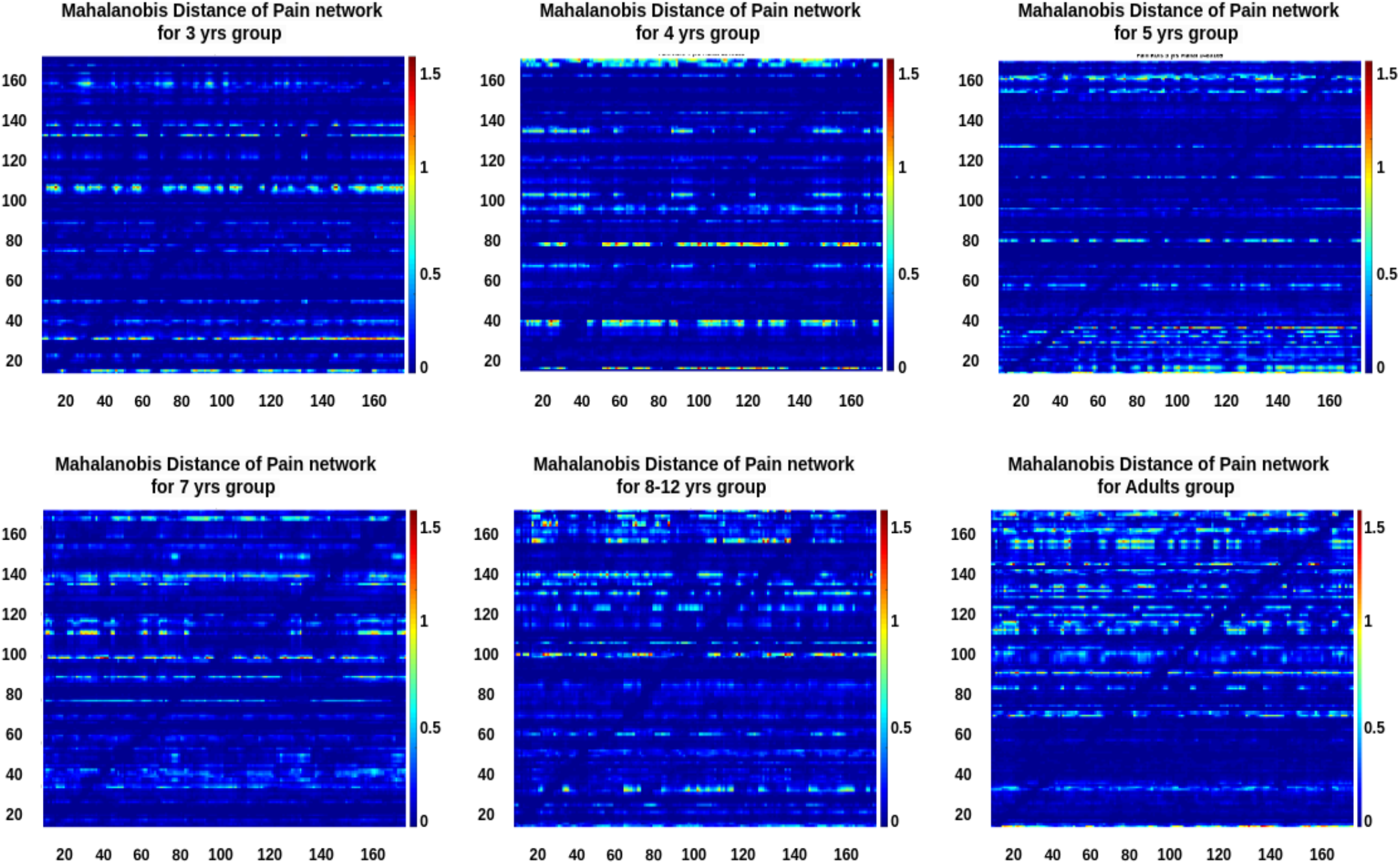
Mahalanobis distance for 6-groups including Pain ROIs

**Figure 4:**
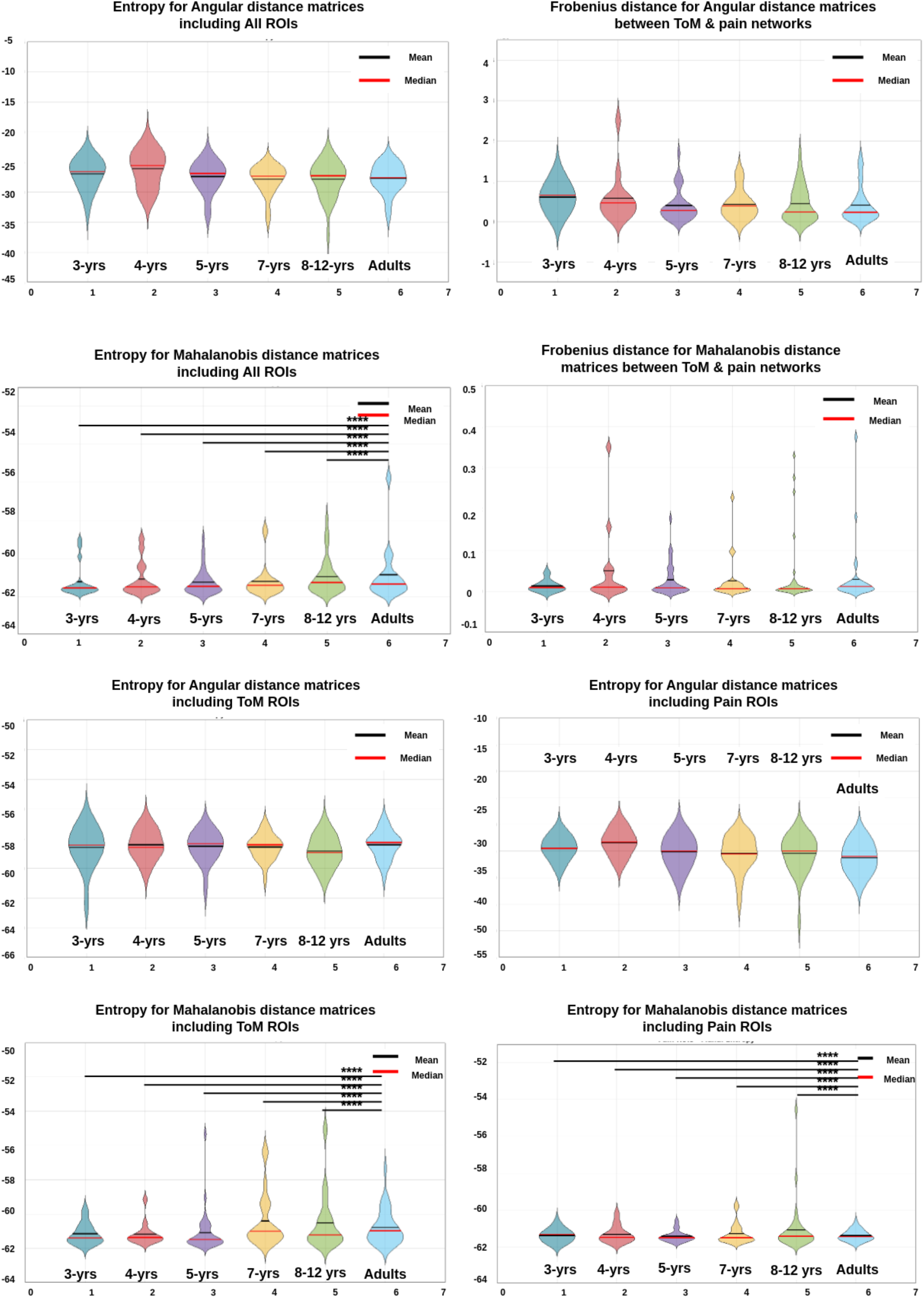
Entropy and Frobenius distances matrices for Angular and Mahalanobis Distances

**Figure 5:**
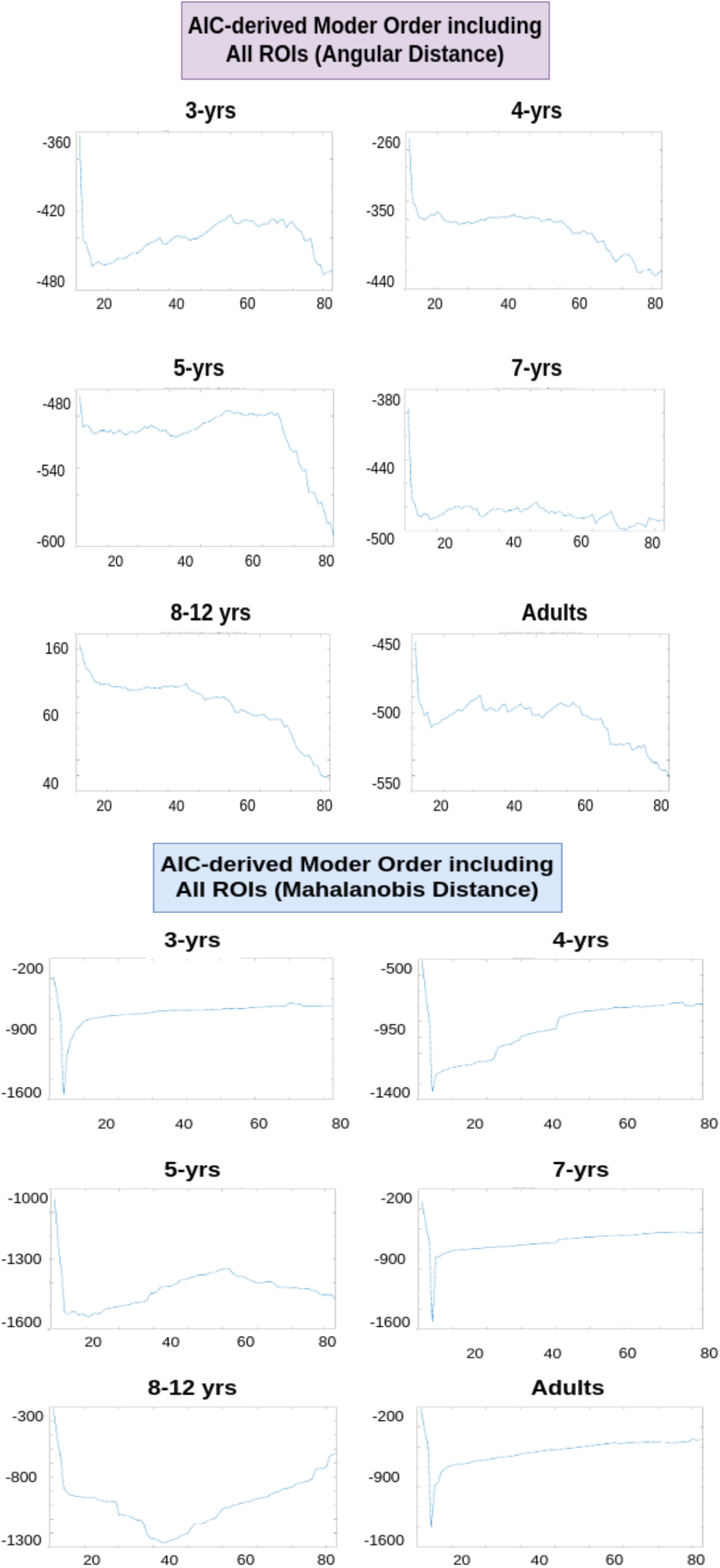
AIC matrices for All ROIs in six-groups analysis, for Angular and Mahalanobis Distances

**Figure 6:**
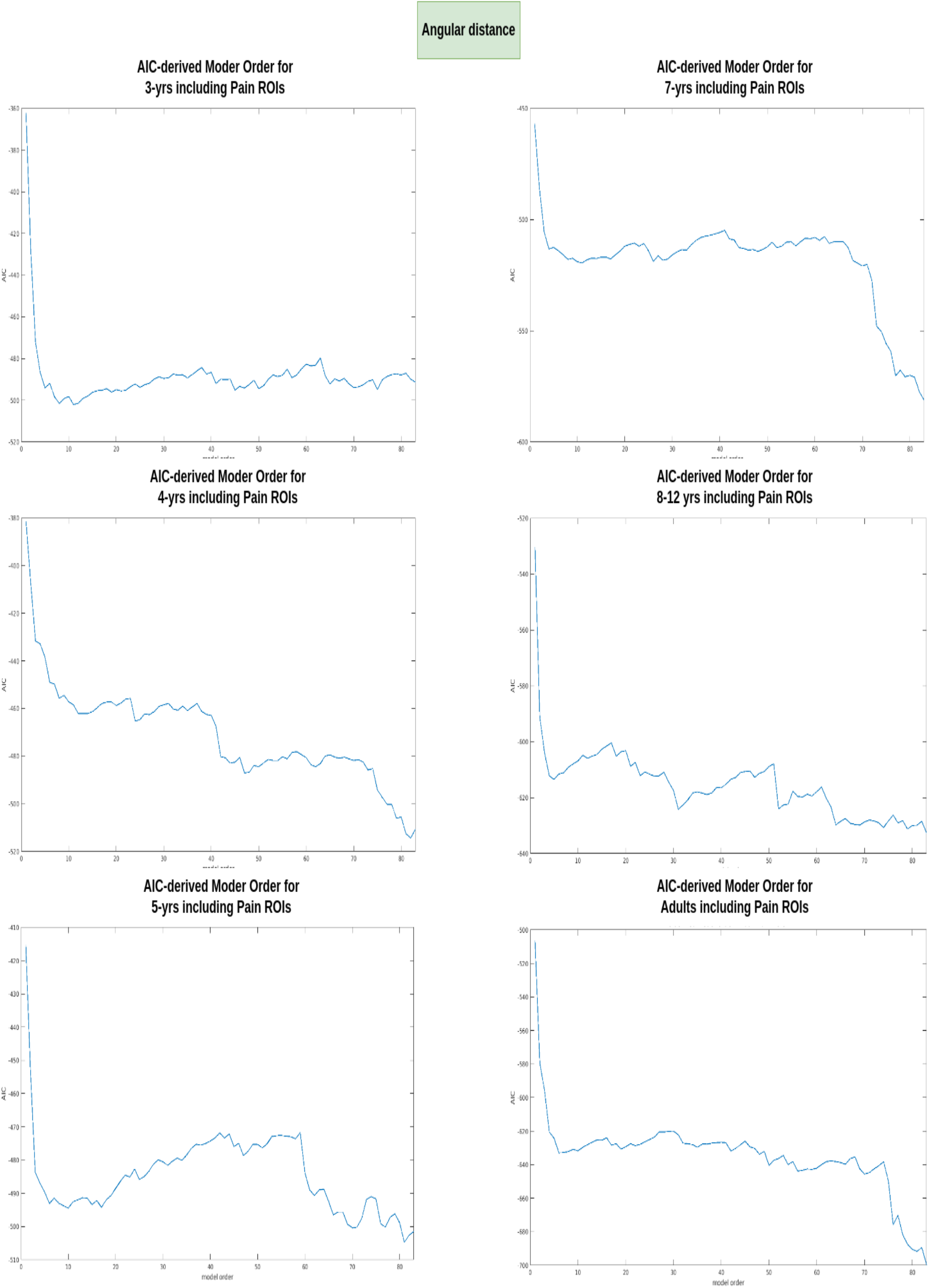
AIC matrices for Pain Network in six-group analysis, for Angular Distance

**Figure 7:**
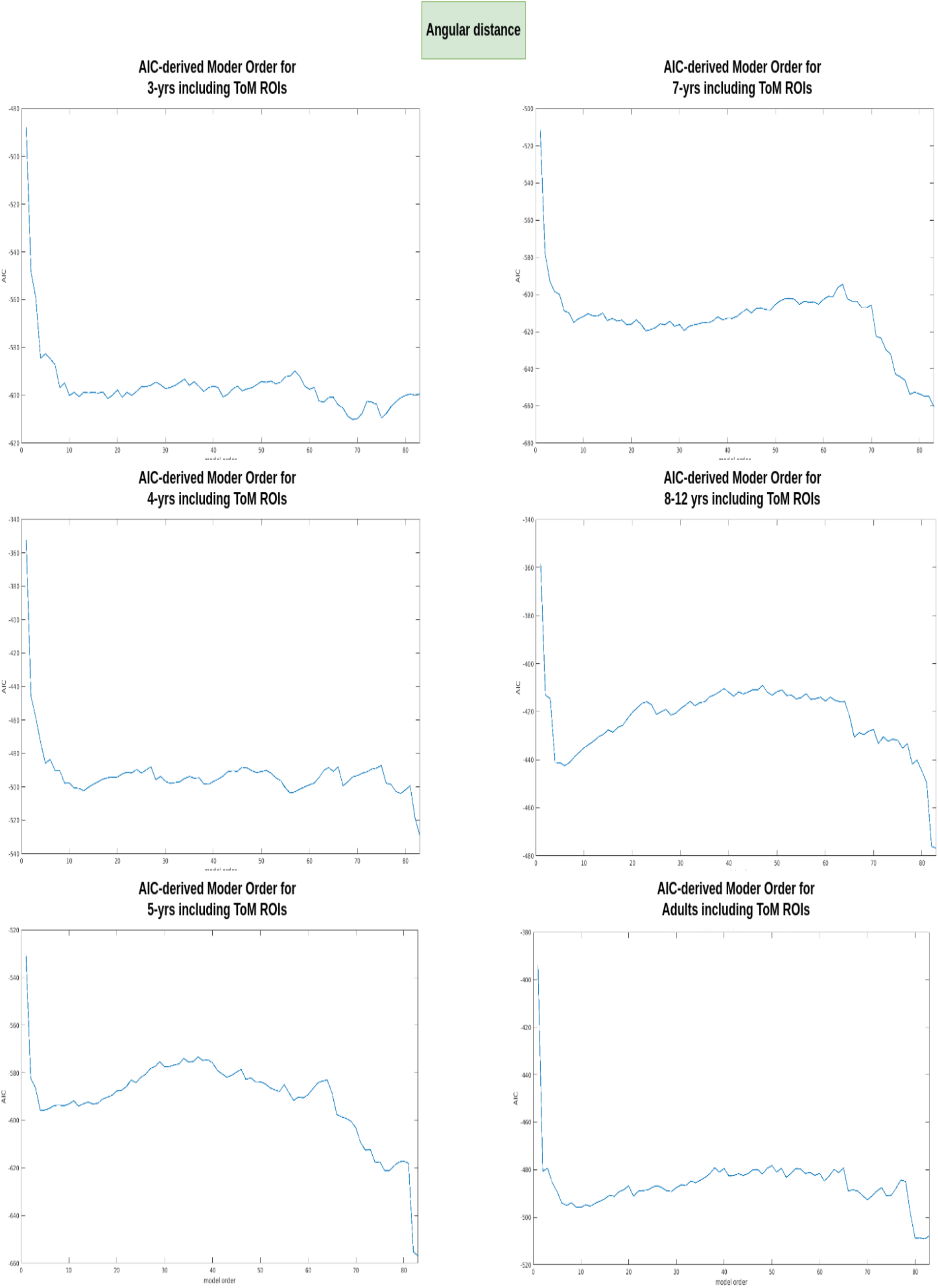
AIC matrices for ToM Network in six-group analysis for Angular Distance

**Figure 8:**
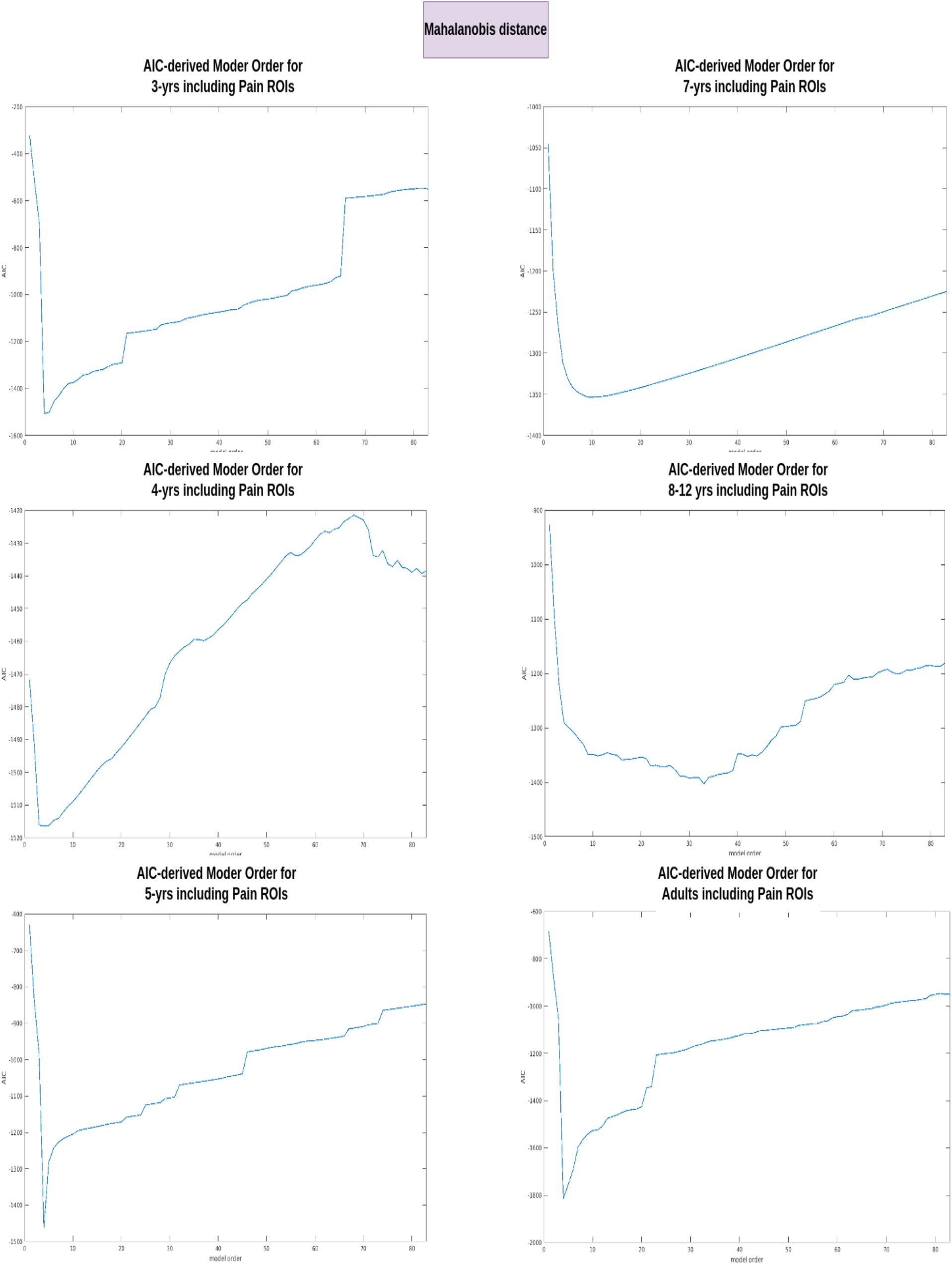
AIC matrices for Pain Network in six-group analysis, for Mahalanobis Distance

**Figure 9:**
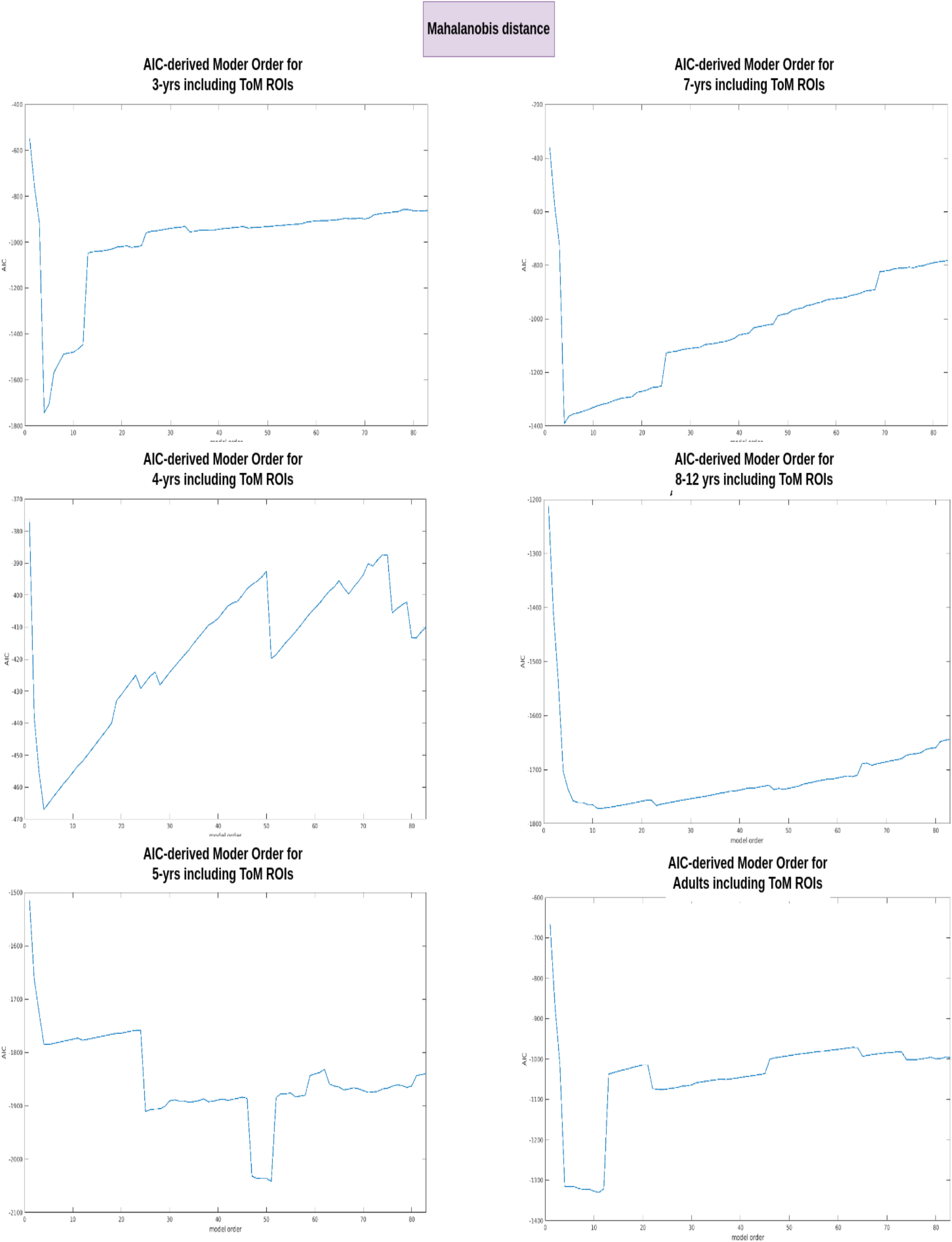
AIC matrices for ToM Network in six-group analysis, for Mahalanobis Distance

#### False-belief Groups Results

**Figure 10:**
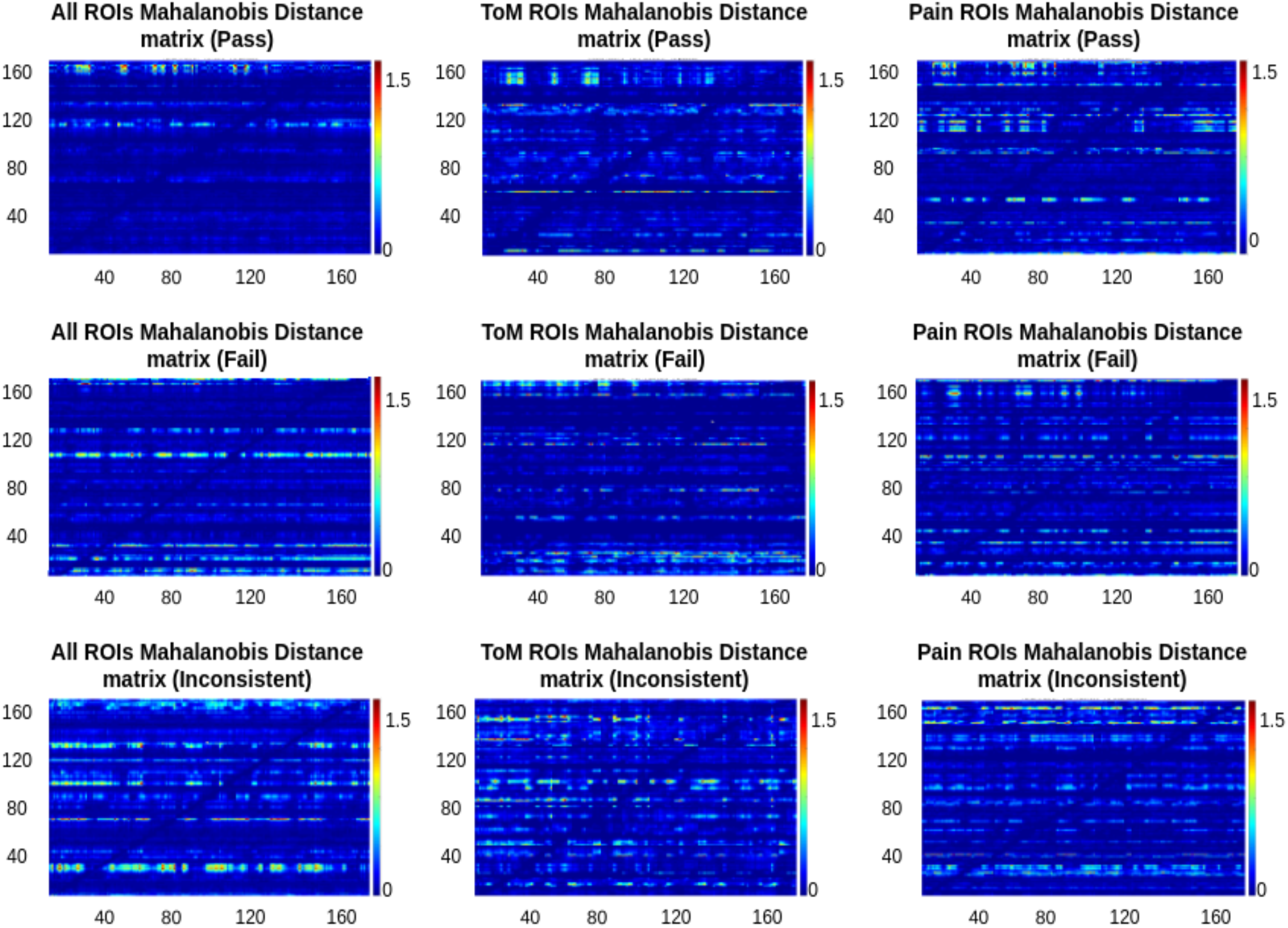
Mahalanobis Distance matrices for pass, fail, & inconsistent groups

**Figure 11:**
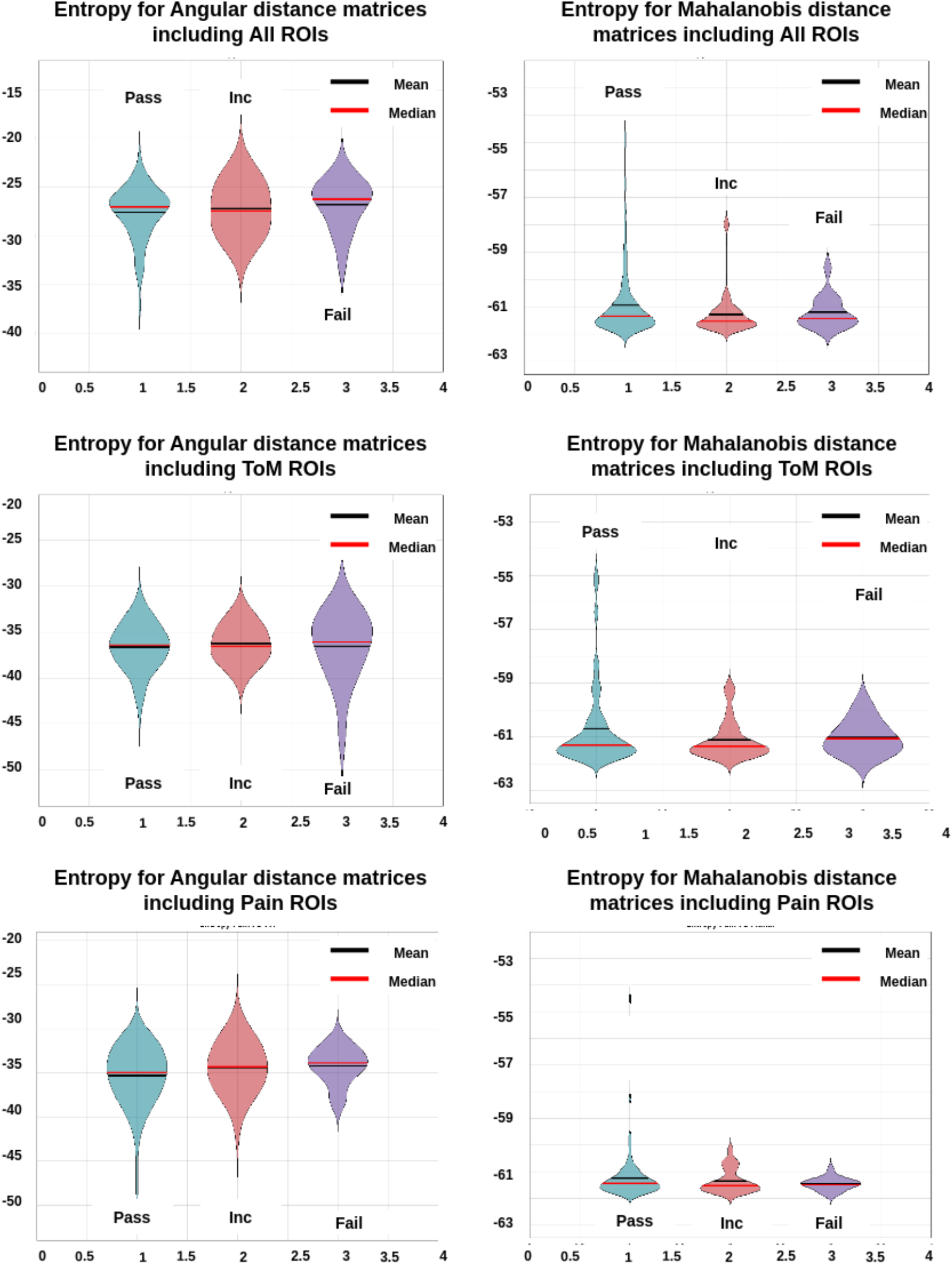
Entropy for pass, fail, & inconsistent groups

**Table 1:**
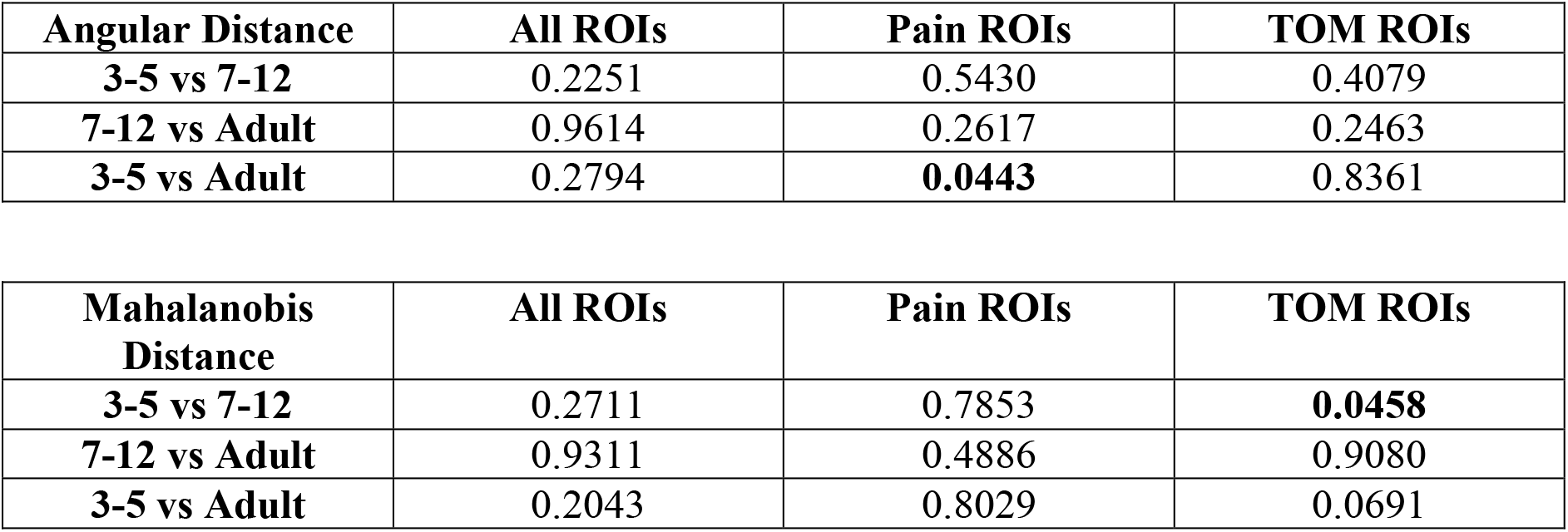
Table with p-values calculated for Kolmogorov-Smirnov test between three (3-5, 7-12 year old, and Adult) age groups of subjects, for Angular and Mahalanobis Distance entropy values.

**Table 2:**
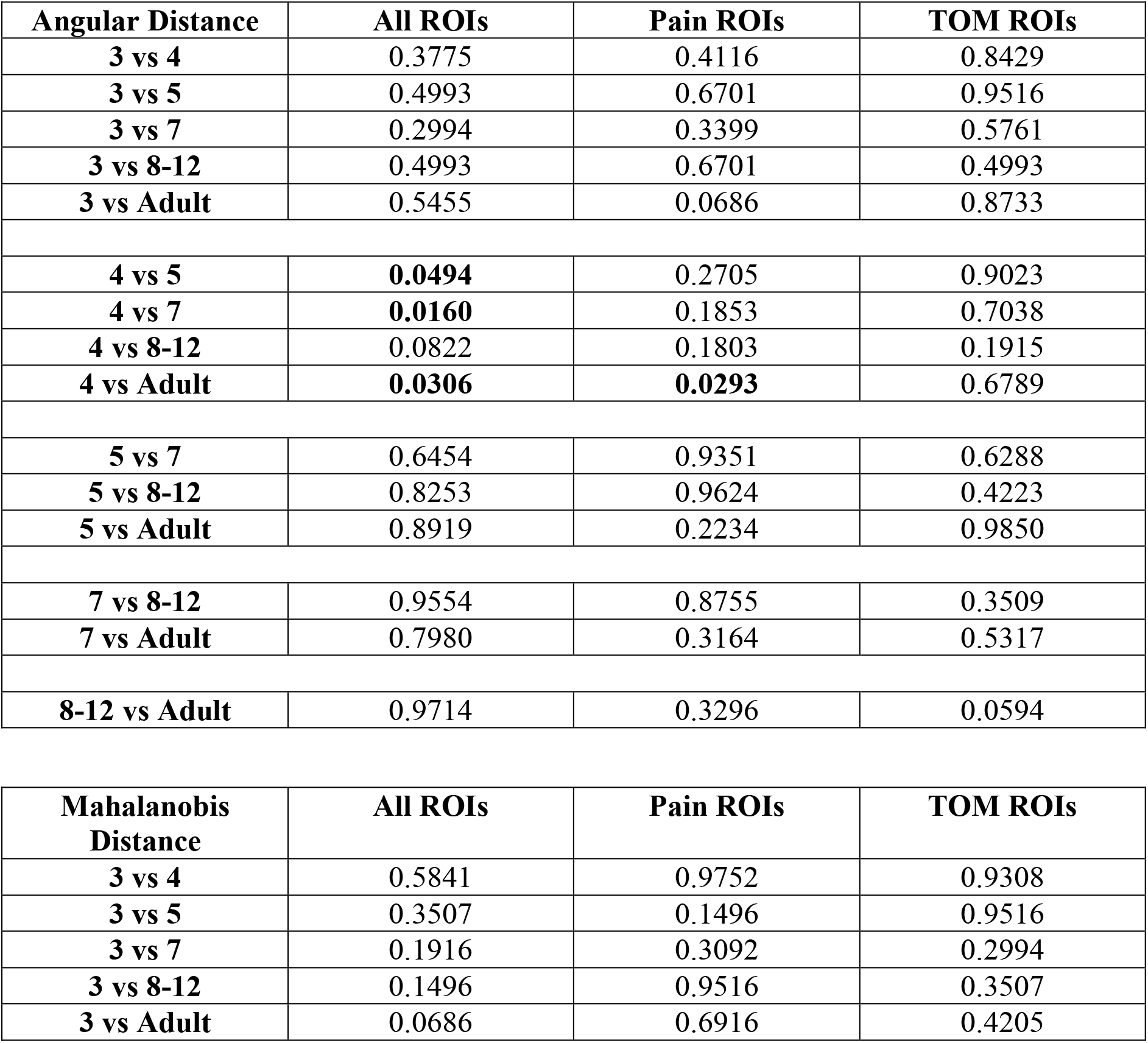
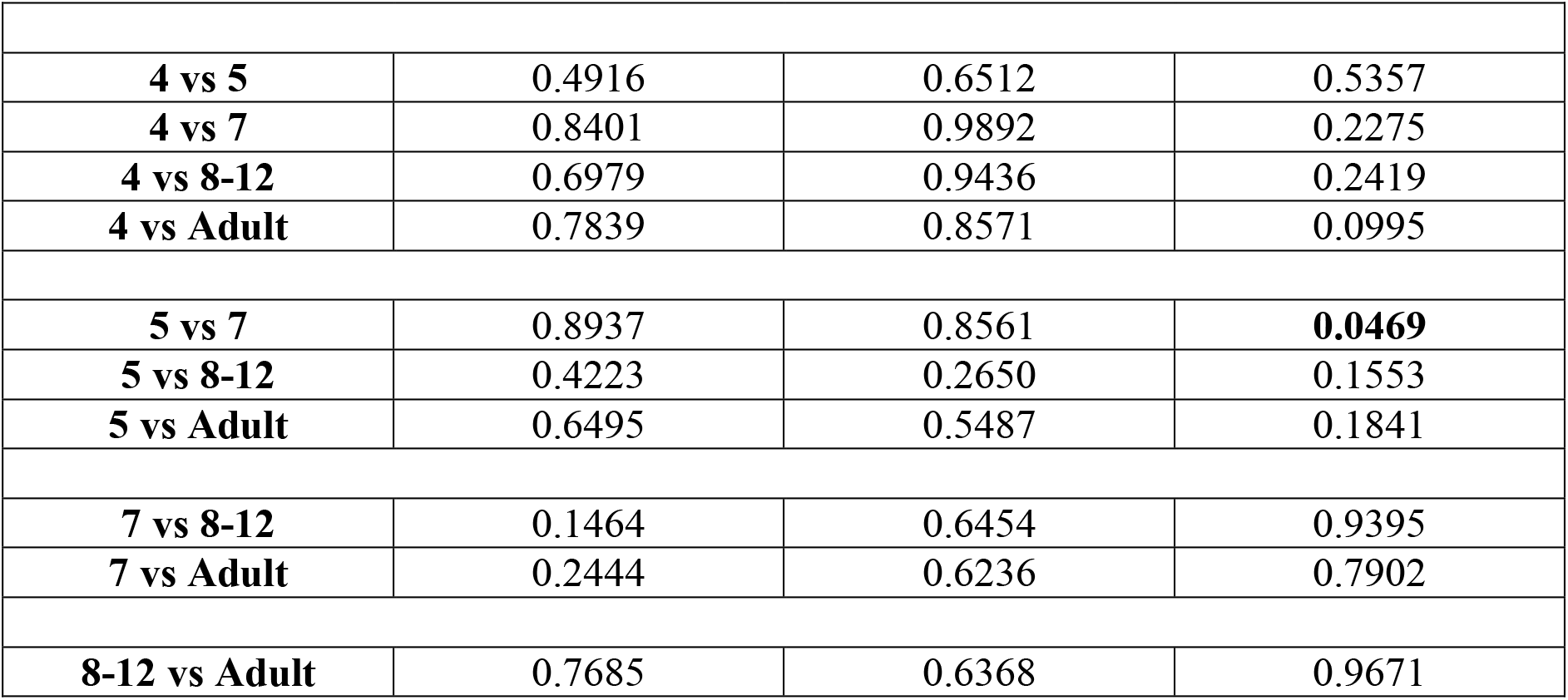
Table with p-values calculated for Kolmogorov-Smirnov test between six (3, 4, 5, 7, 8-12 years old, and Adult) age groups of subjects, for Angular and Mahalanobis Distance entropy values.

**Table 3:**
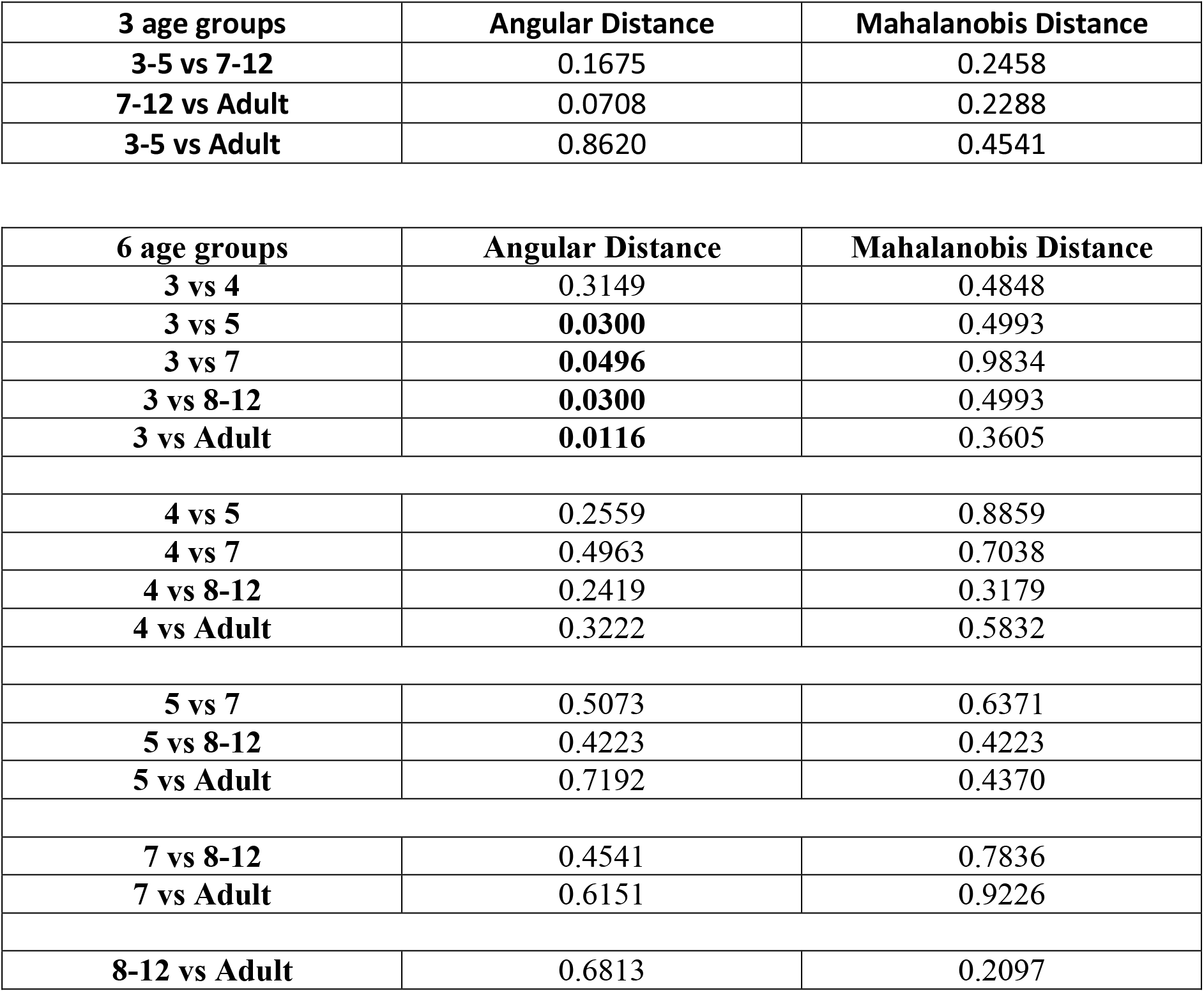
Table with p-values calculated for Kolmogorov-Smirnov test for three (3-5, 7-12-year-old, and Adult) and six (3, 4, 5, 7, 8-12, and Adult) age group subjects, Frobenius Distance between TOM and Pain Angular and Mahalanobis Distances.

**Table 4:**
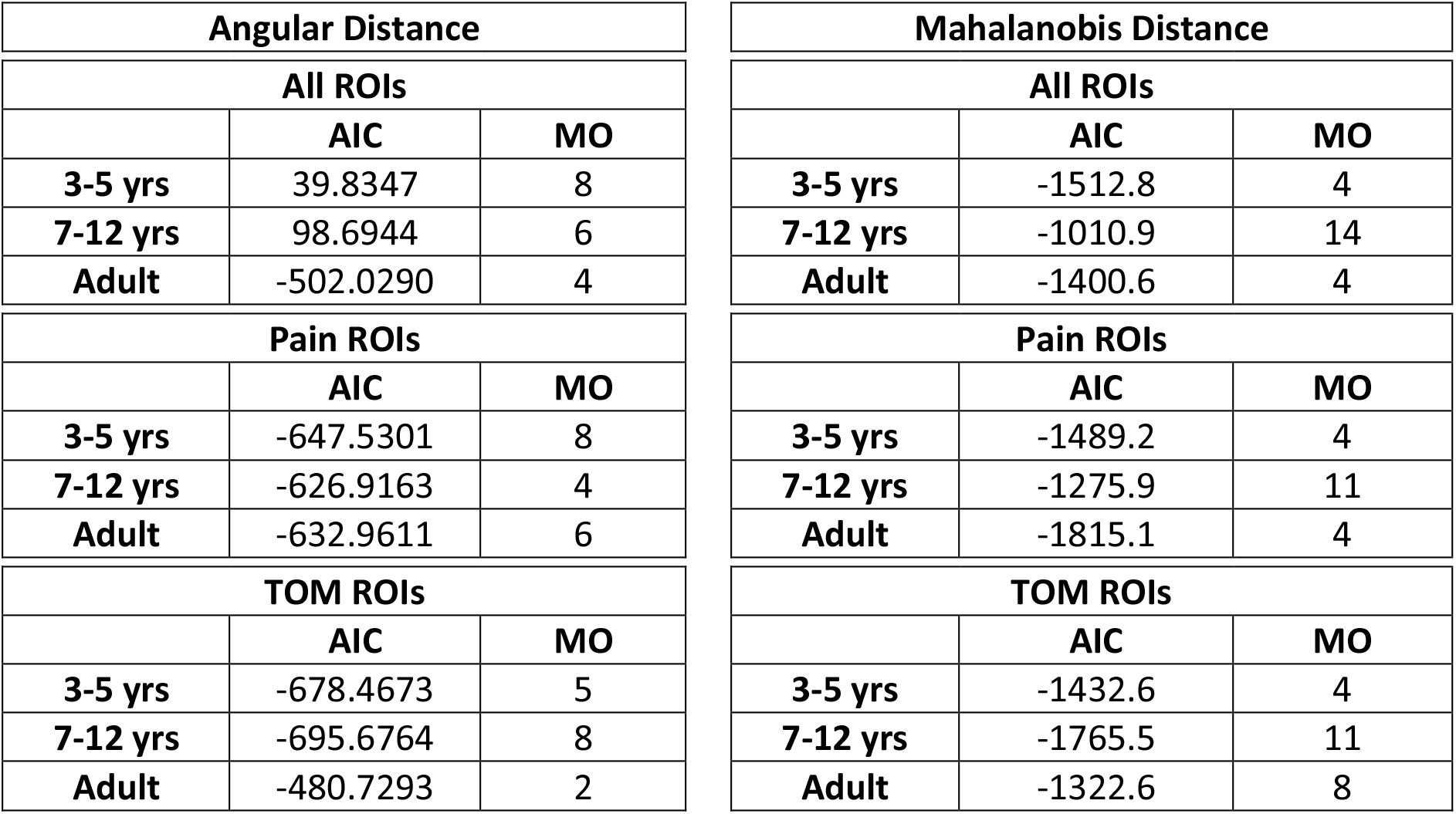
Table with (the first) minimal AIC score and the corresponding Model Order number, obtained for the stochastic characterization of the Angular and Mahalanobis Distance temporal matrices for three age group (3-5, 7-12 year olds and Adult) subjects.

**Table 5:**
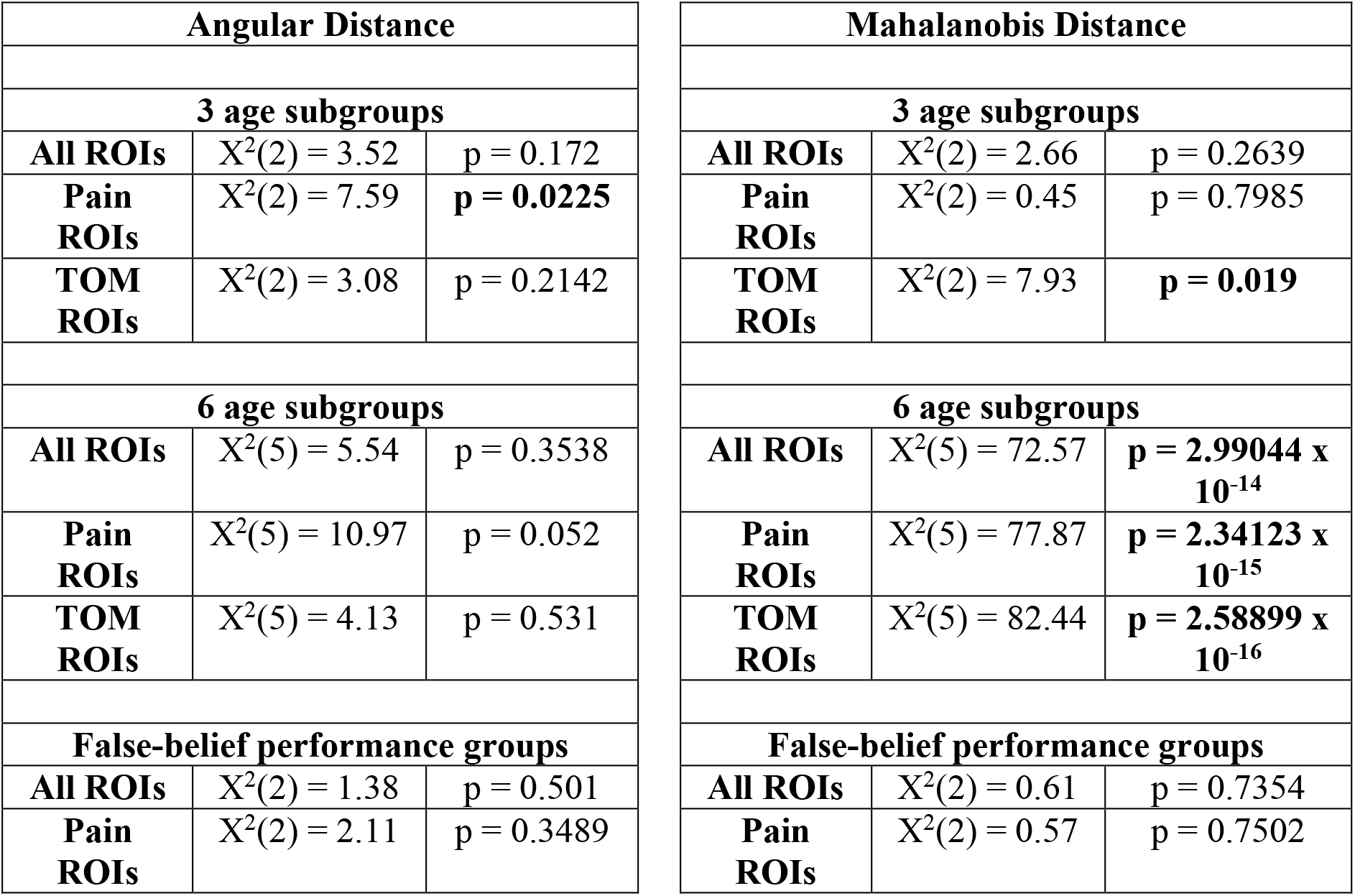
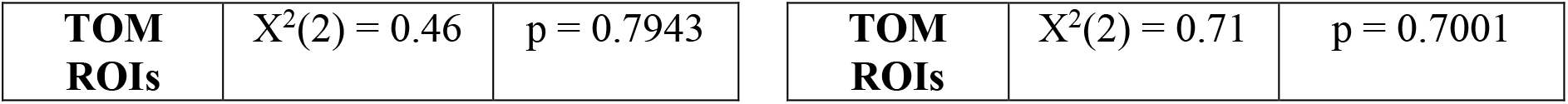
Table with values calculated for Kruskal-Wallis test for three (3-5, 7-12 yrs and Adult), six (3, 4, 5, 7, 8-12 yrs and Adult) age groups and False-belief performance (Pass, Inconsistent, Fail) groups for Angular and Mahalanobis Distance-derived Entropy.

**Table 6:**
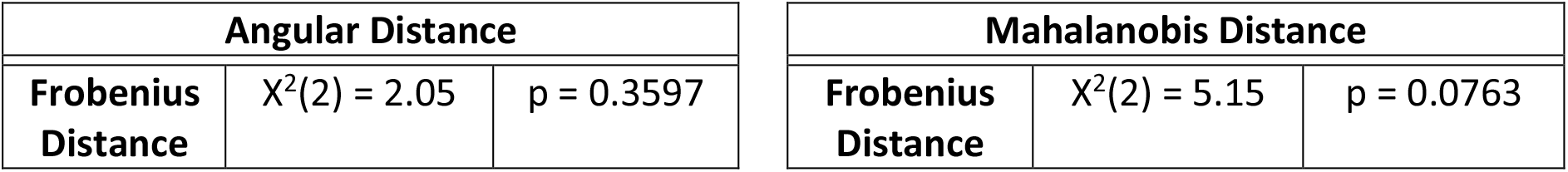
Table with values calculated for Kruskal-Wallis test for three (3-5, 7-12 yrs and Adult) age groups for Angular Distance and Mahalanobis Distance-derived Frobenius Distance.

**Table 7:**
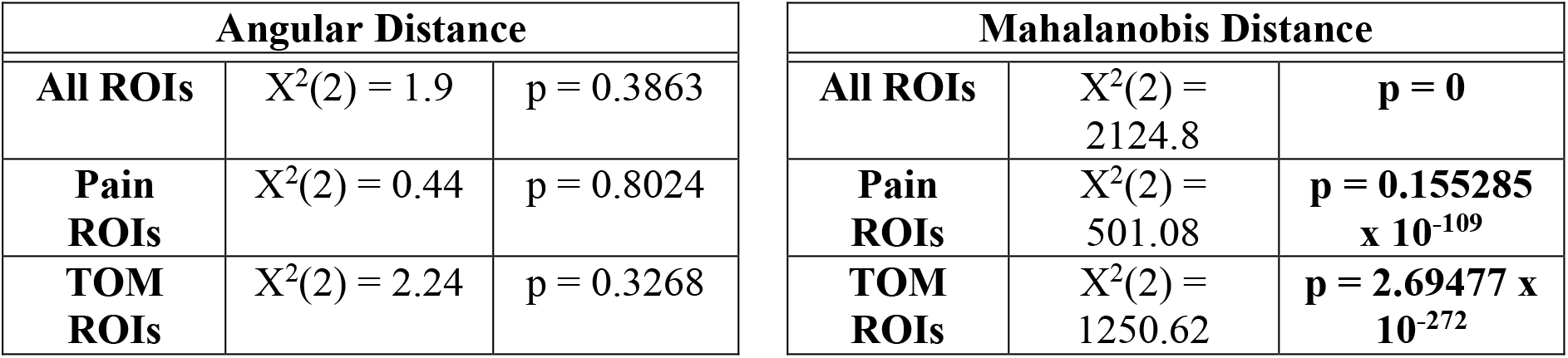
Table with values calculated for Kruskal-Wallis test for three (3-5, 7-12 yrs and Adult) age groups for Angular Distance and Mahalanobis Distance-derived Z-scores.

**Table 8:**
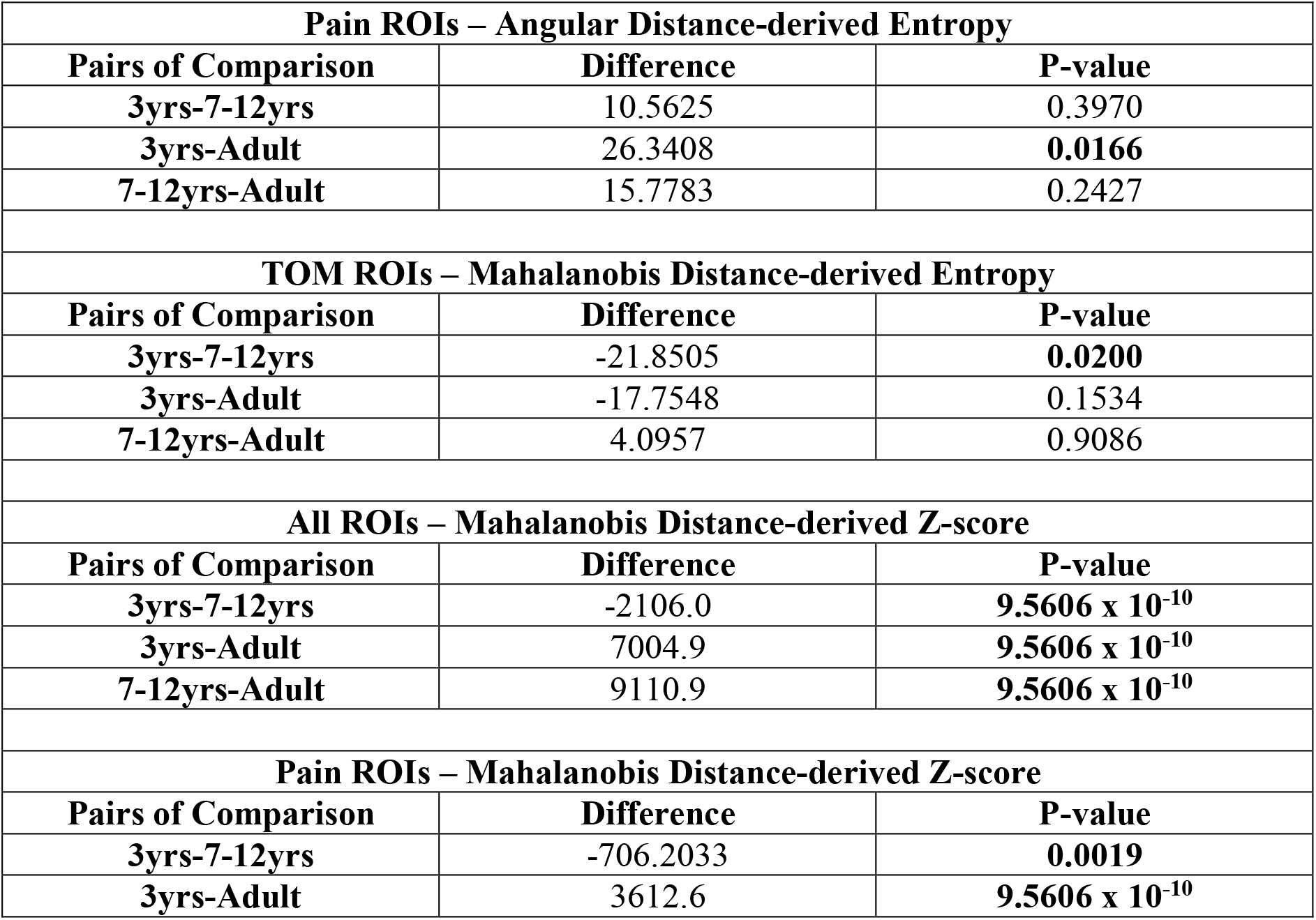
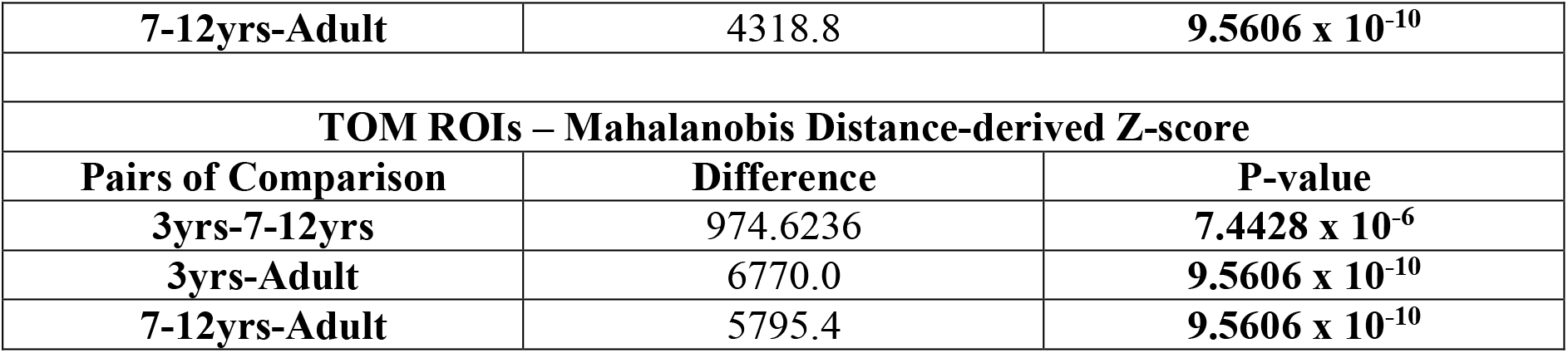
Table with values calculated for Dunn-Sidak post-hoc test for significant values in three (3-5, 7-12 yrs and Adult) age groups.

**Table 8:**
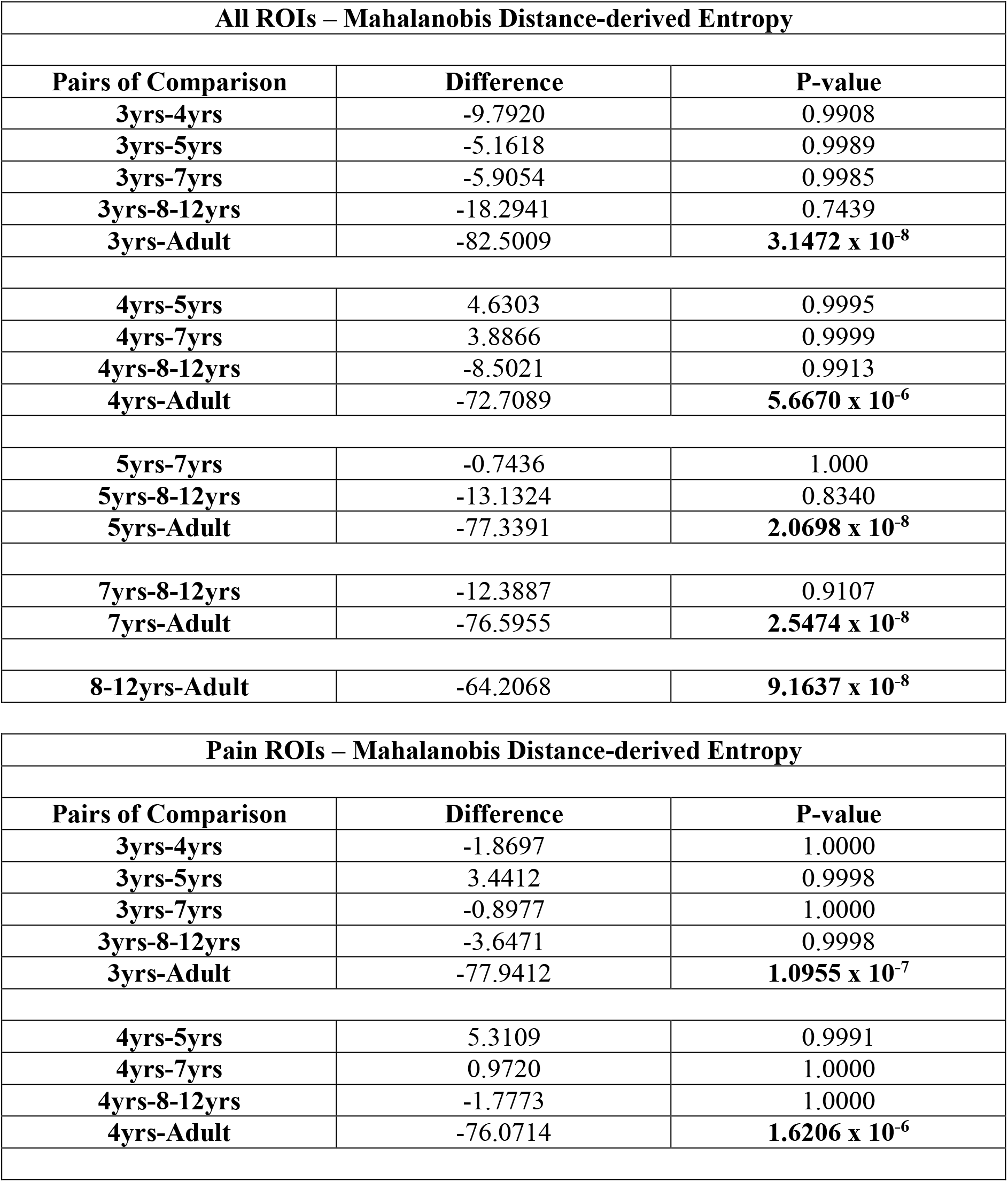
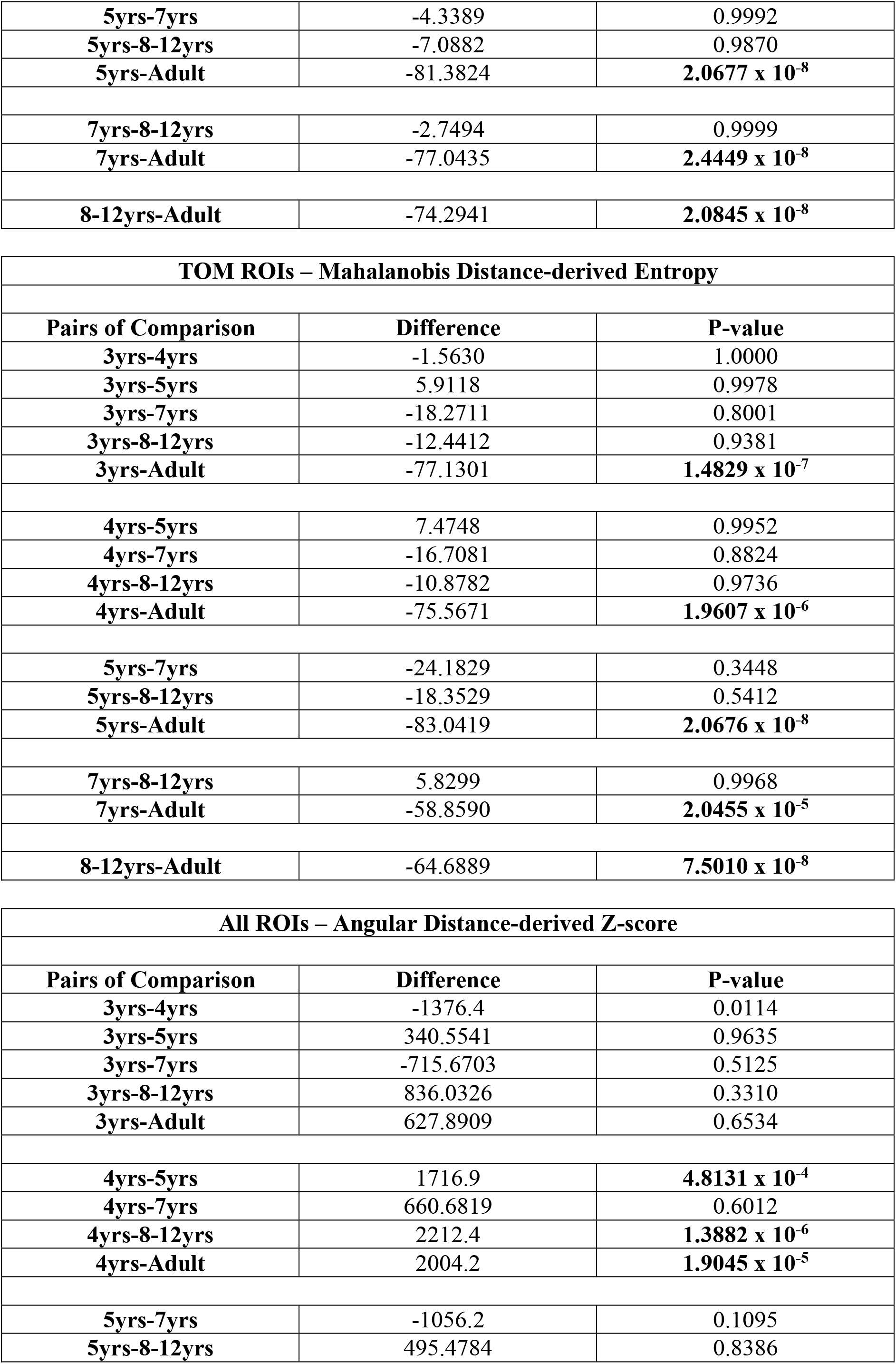
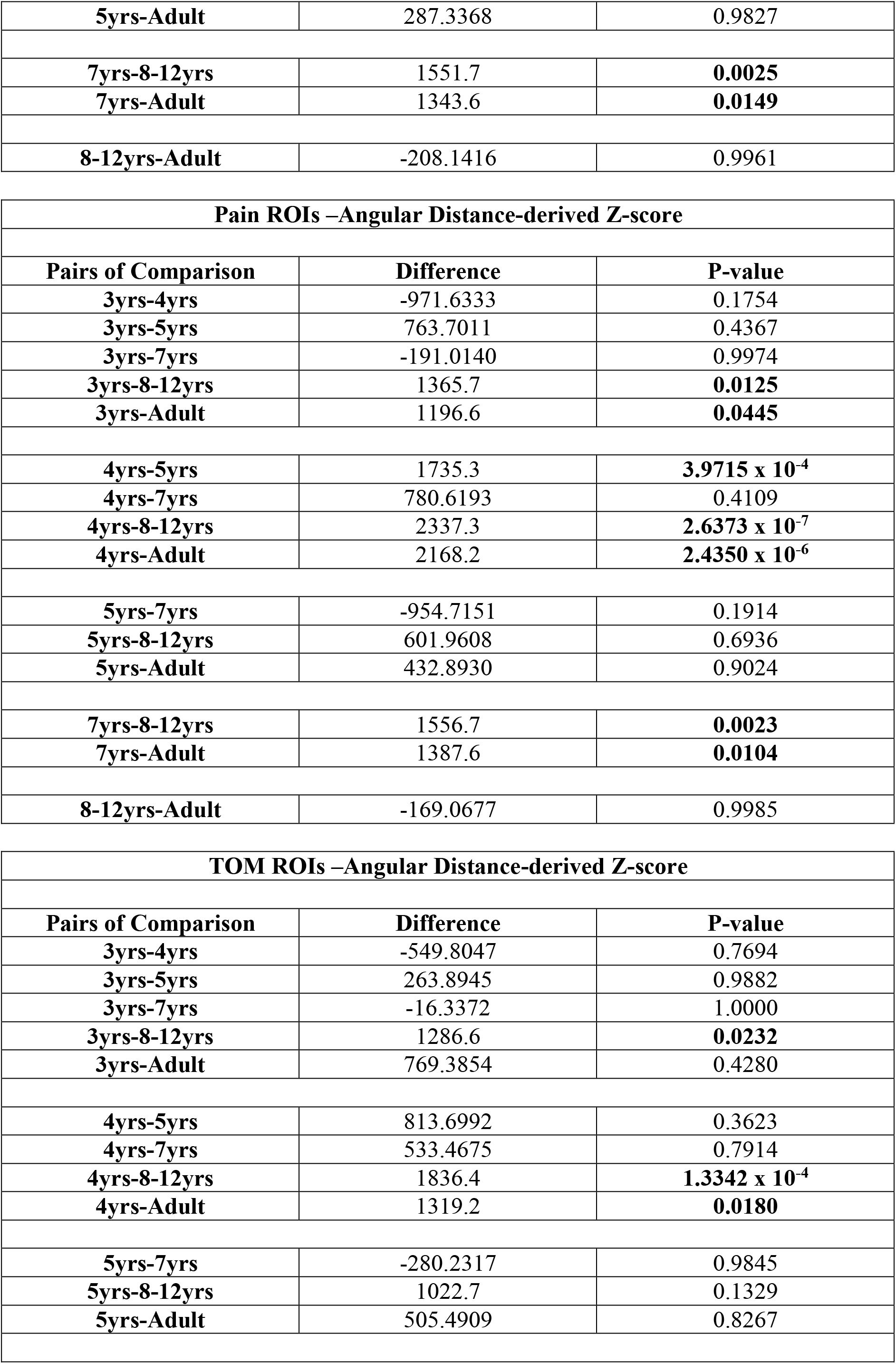
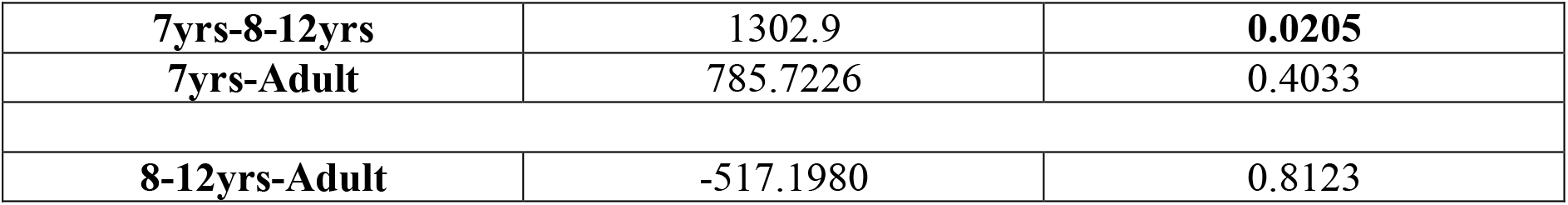
Table with values calculated for Dunn-Sidak post-hoc test for significant values in six (3, 4, 5, 7, 8-12 yrs and Adult) age groups.

**Table 9:** Table with (the first) minimal AIC score and the corresponding Model Order number, obtained for the stochastic characterization of the Angular and Mahalanobis Distance temporal matrices for 6 age groups (3, 4, 5, 7, 8-12, and Adult) of subjects.

| **Angular Distance** |  |  | **Mahalanobis Distance** |  |  |
| --- | --- | --- | --- | --- | --- |
| **All ROIs** |  |  | **All ROIs** |  |  |
|  | **AIC** | **MO** |  | **AIC** | **MO** |
| **3 yrs** | -461.39 | 5 | **3 yrs** | -1550.3 | 4 |
| **4 yrs** | -360.21 | 6 | **4 yrs** | -1349.6 | 4 |
| **5 yrs** | -508.79 | 2 | **5 yrs** | -1535.1 | 5 |
| **7 yrs** | -488.25 | 5 | **7 yrs** | -1525.4 | 4 |
| **8-12 yrs** | 98.69 | 6 | **8-12 yrs** | -944.94 | 11 |
| **Adult** | -502.03 | 4 | **Adult** | -1440.6 | 4 |
| **Pain ROIs** |  |  | **Pain ROIs** |  |  |
|  | **AIC** | **MO** |  | **AIC** | **MO** |
| **3 yrs** | -494.14 | 5 | **3 yrs** | -1506.8 | 4 |
| **4 yrs** | -455.69 | 8 | **4 yrs** | -1516.4 | 4 |
| **5 yrs** | -493.04 | 6 | **5 yrs** | -1461.9 | 4 |
| **7 yrs** | -513.25 | 4 | **7 yrs** | -1353.5 | 10 |
| **8-12 yrs** | -613.47 | 5 | **8-12 yrs** | -1349.4 | 4 |
| **Adult** | -632.96 | 6 | **Adult** | -1815.1 | 4 |
| **ToM ROIs** |  |  | **ToM ROIs** |  |  |
|  | **AIC** | **MO** |  | **AIC** | **MO** |
| **3 yrs** | -584.51 | 4 | **3 yrs** | -1743.5 | 4 |
| **4 yrs** | -485.89 | 5 | **4 yrs** | -467.01 | 4 |
| **5 yrs** | -595.74 | 4 | **5 yrs** | -1785.2 | 5 |
| **7 yrs** | -615.09 | 8 | **7 yrs** | -1391.5 | 4 |
| **8-12 yrs** | -442.53 | 6 | **8-12 yrs** | -1771.7 | 11 |
| **Adult** | -480.73 | 2 | **Adult** | -1322.6 | 8 |

#### Results for ISC

**Figure 12:**
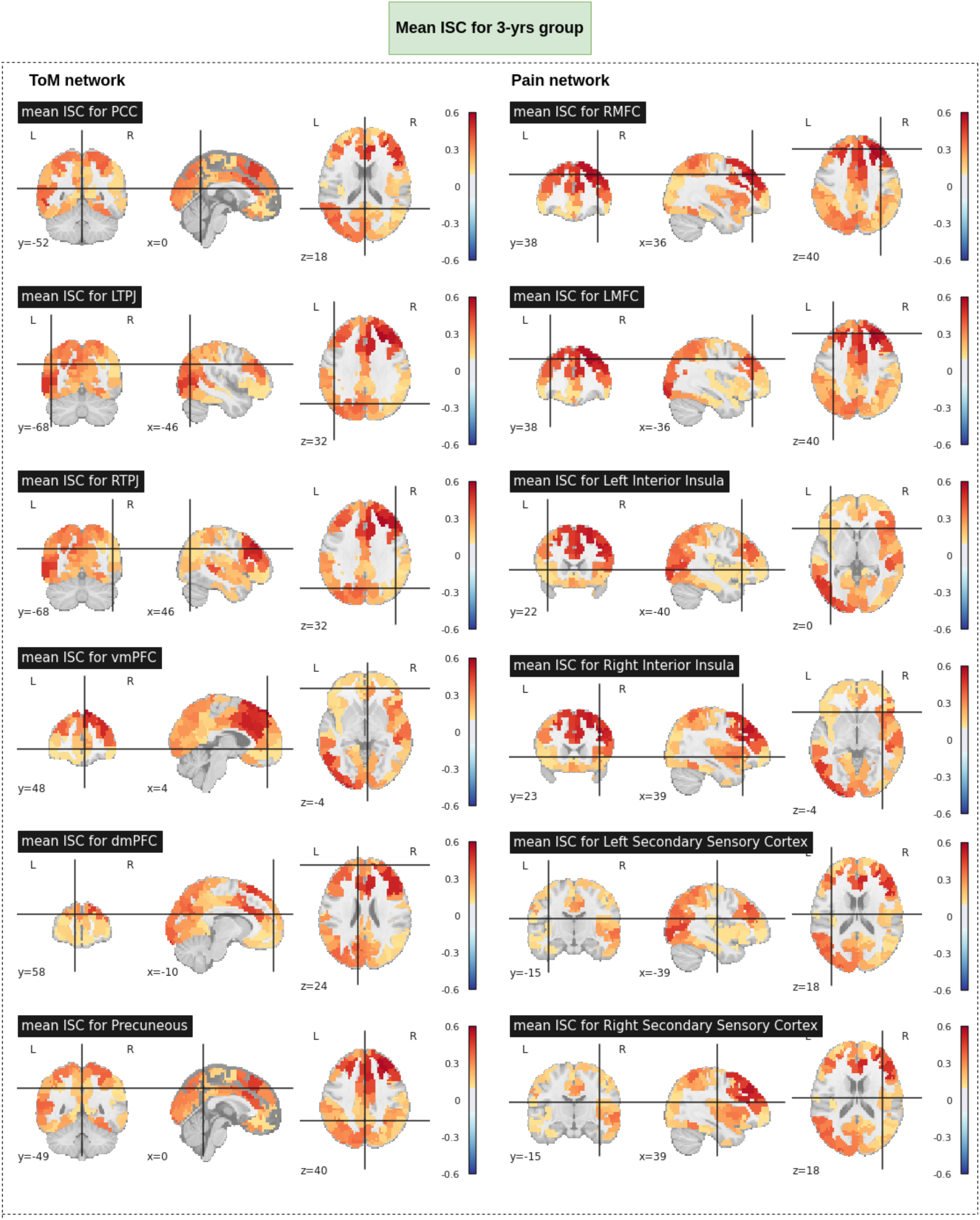
Mean ISC for 3-yrs age group

**Figure 13:**
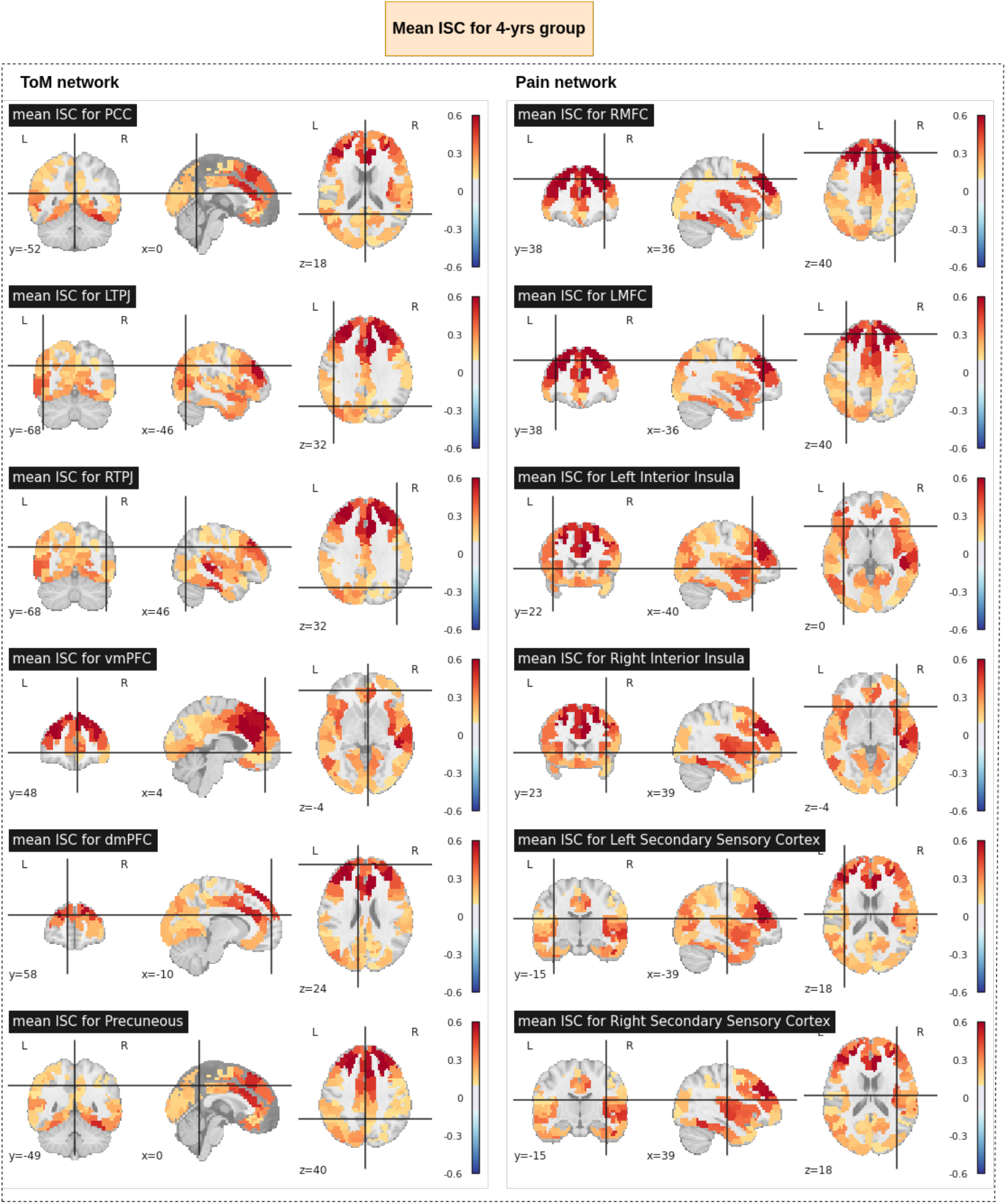
Mean ISC for 4-yrs age group

**Figure 14:**
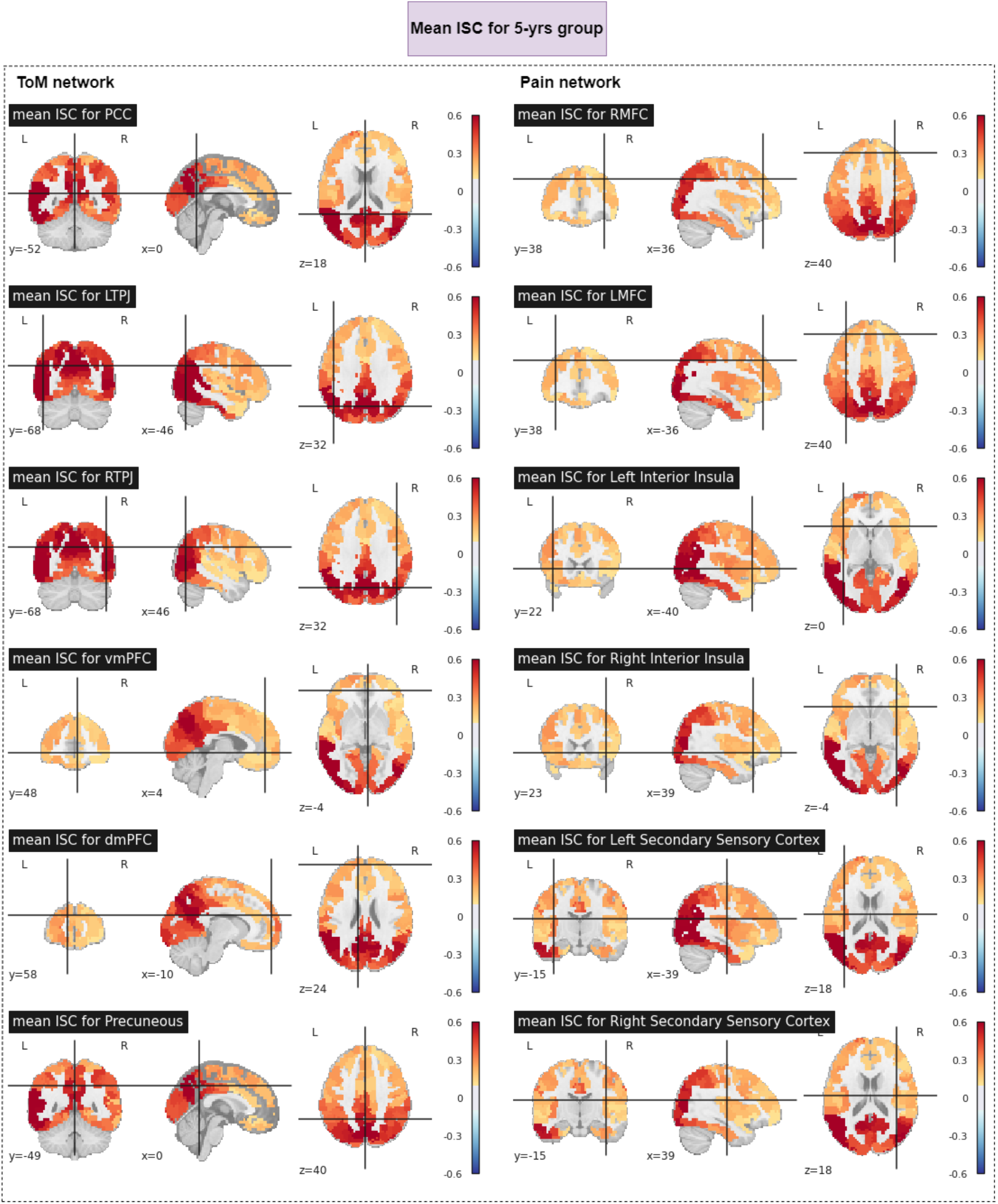
Mean ISC for 5-yrs age group

**Figure 15:**
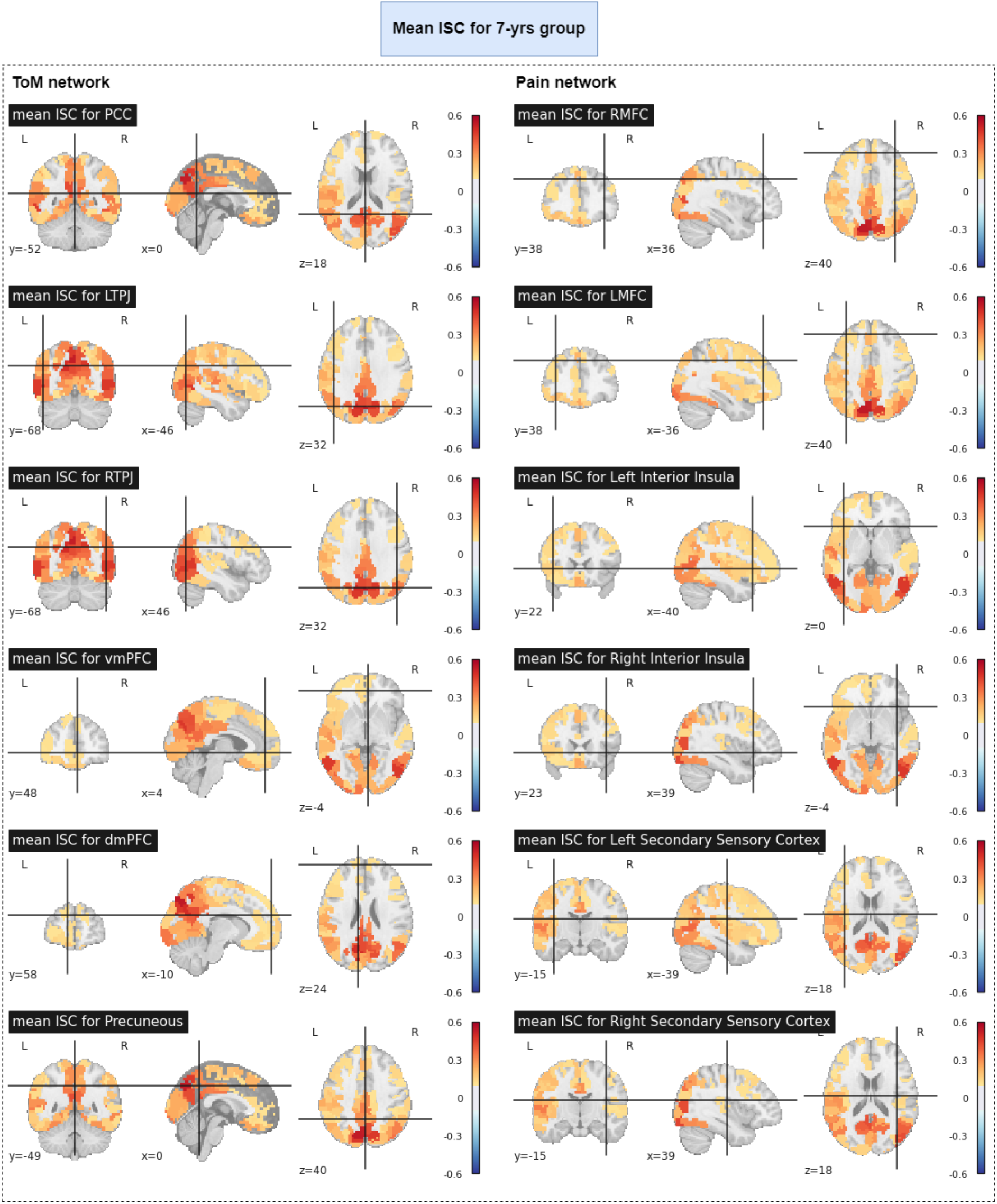
Mean ISC for 7-yrs age group

**Figure 16:**
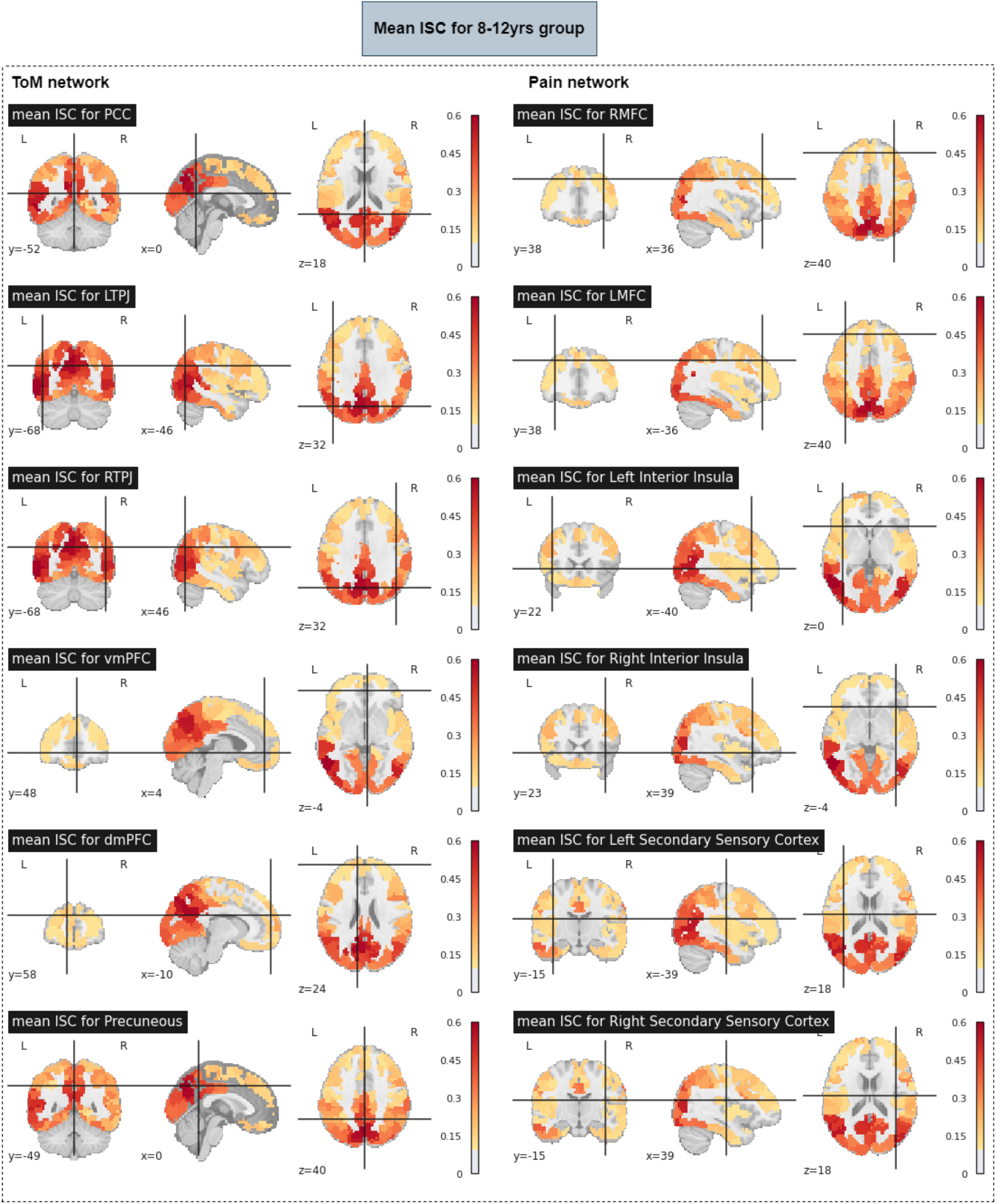
Mean ISC for 8-12 yrs age group

**Figure 17:**
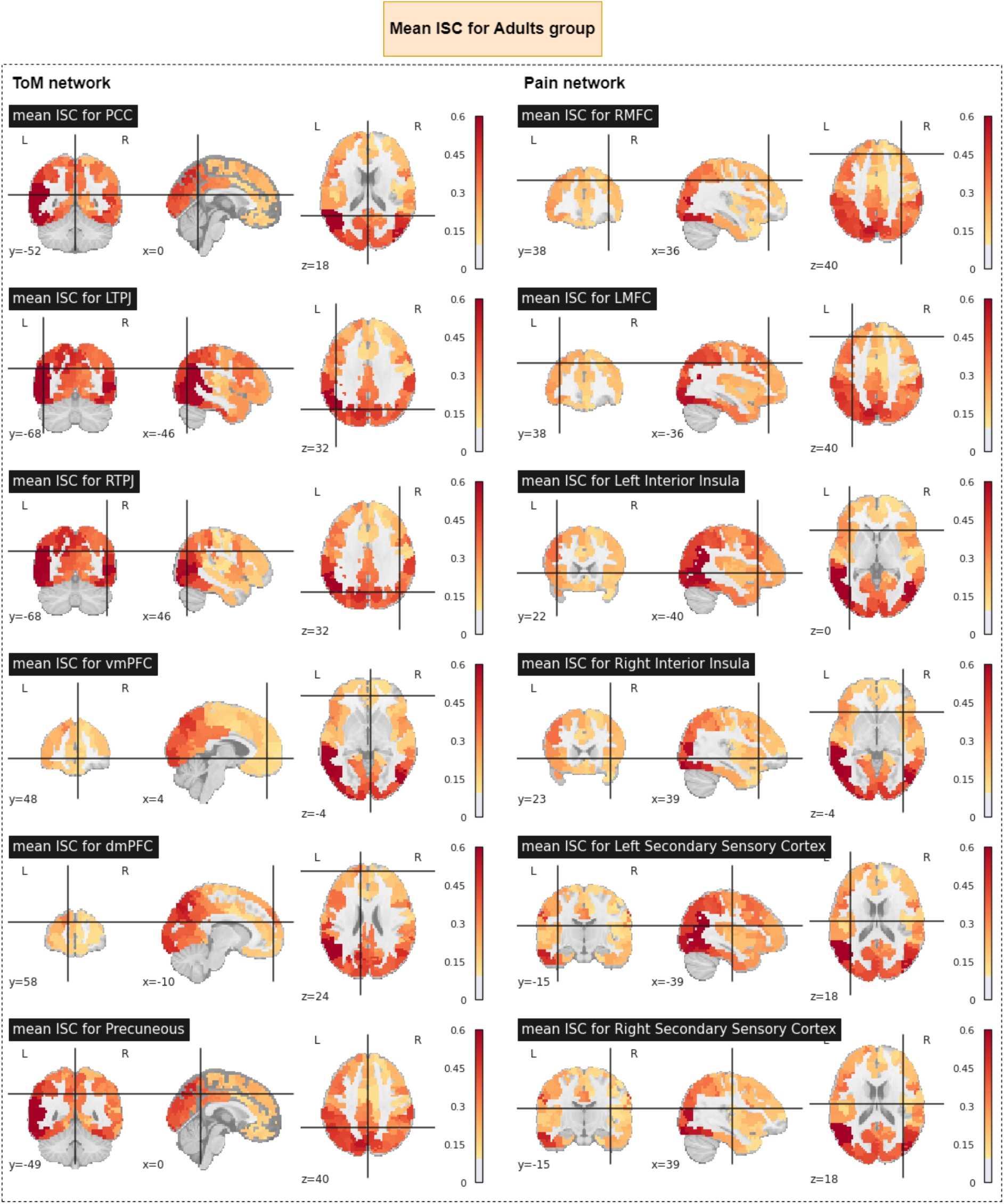
Mean ISC for Adult Group

**Figure 18:**
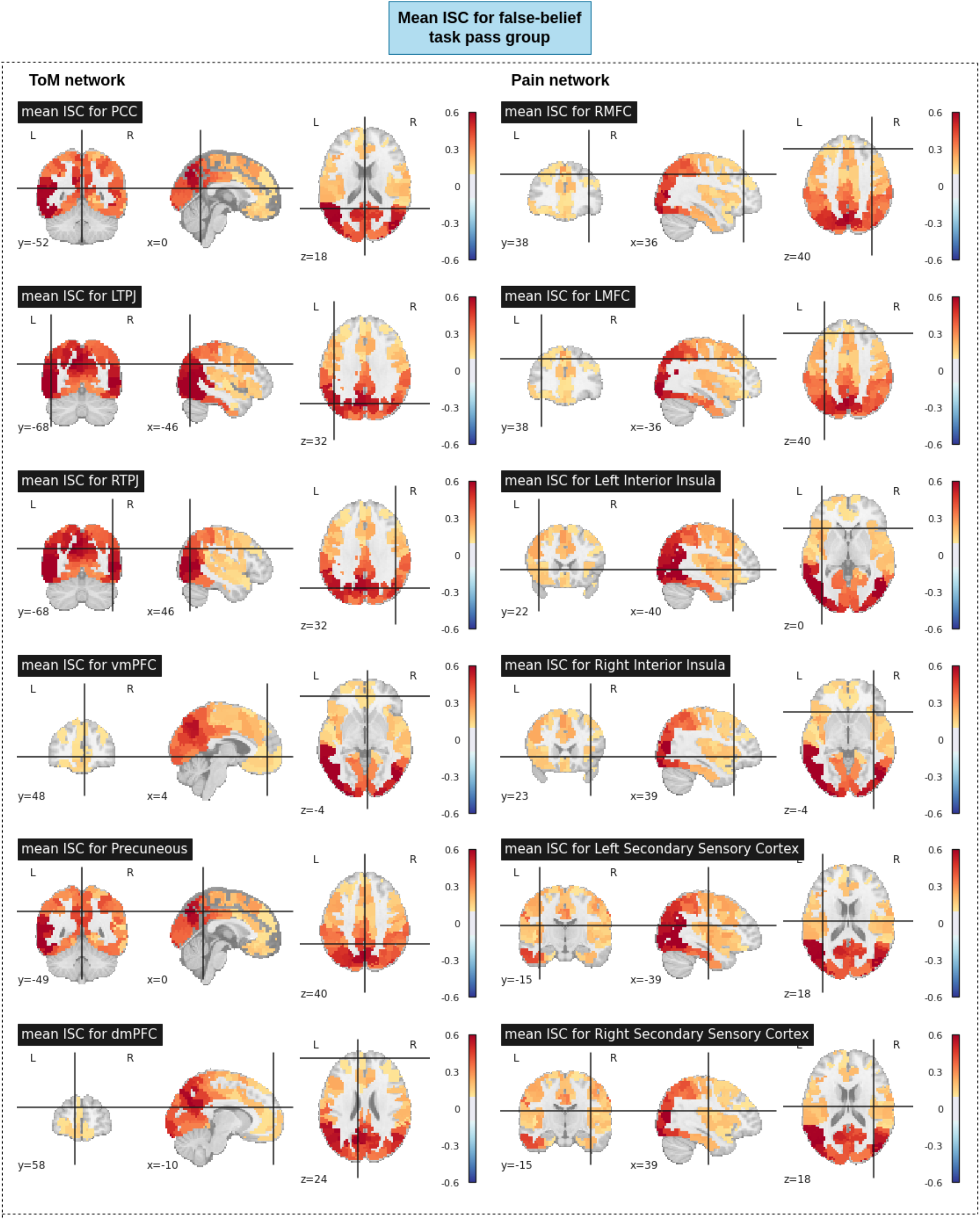
Mean ISC for false-belief task pass group

**Figure 19:**
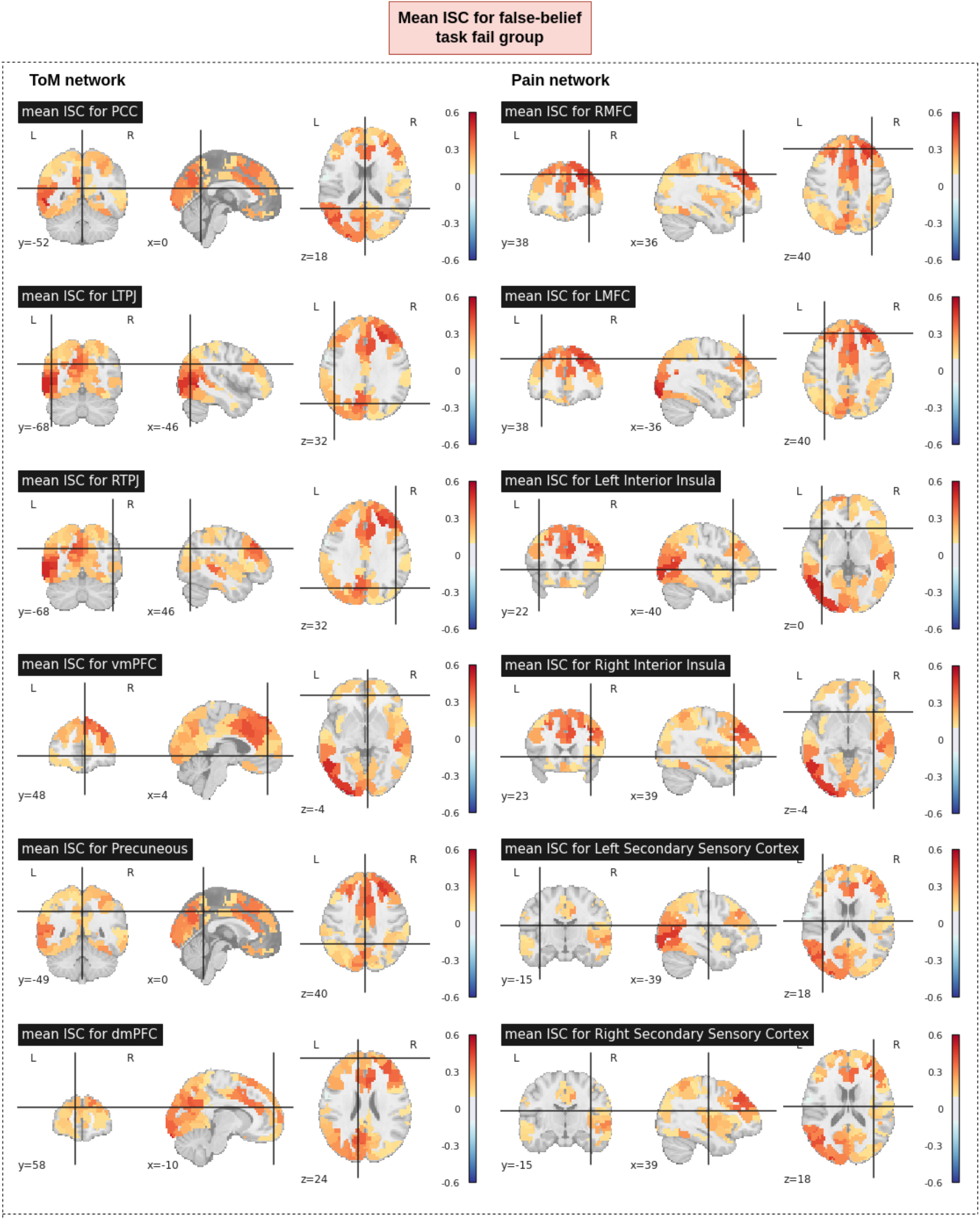
Mean ISC for false-belief task fail group

**Figure 20:**
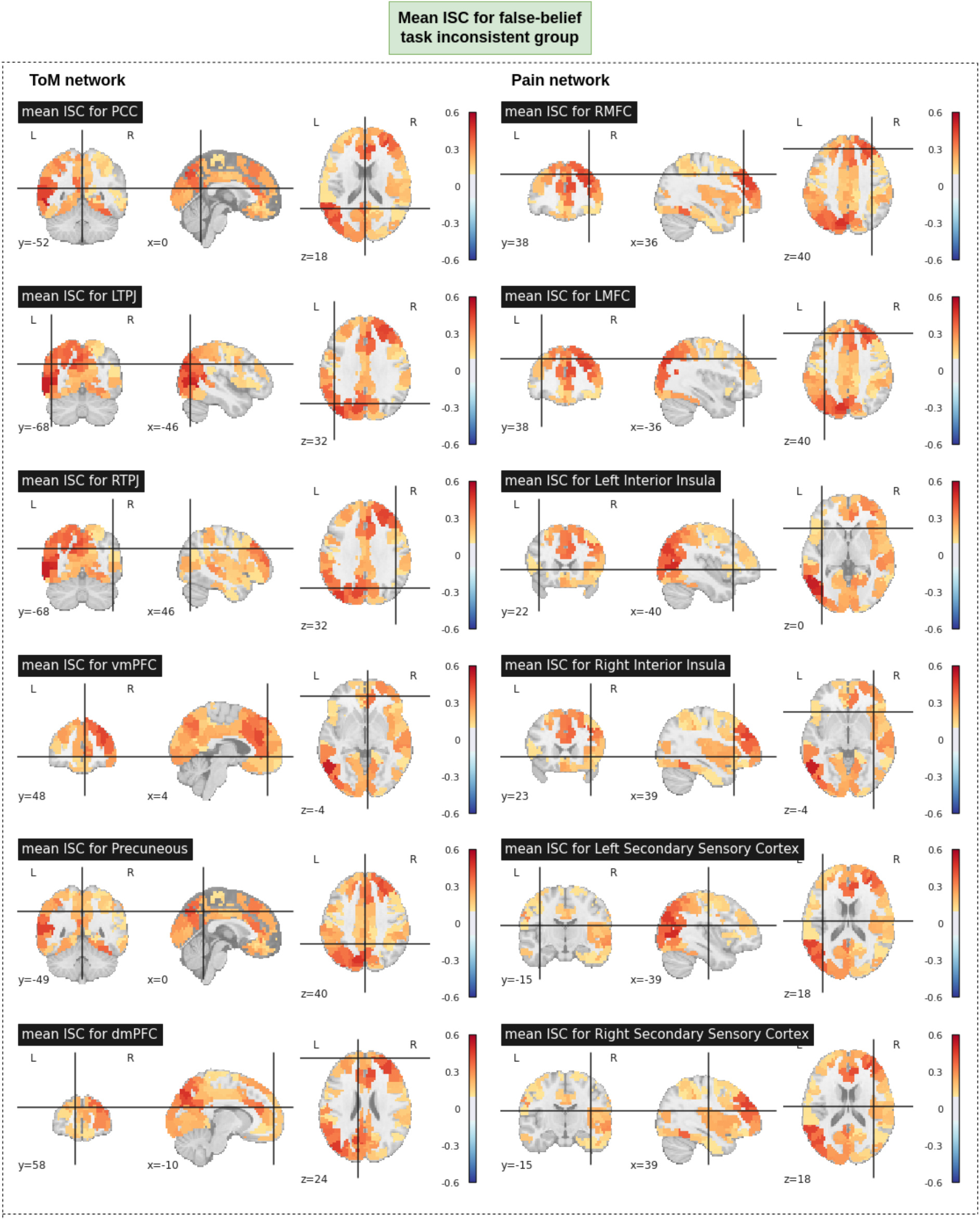
Mean ISC for false-belief task inconsistent group

